# A Human Brain-Chip for modelling brain pathologies and screening of BBB crossing therapeutic strategies

**DOI:** 10.1101/2024.08.11.607509

**Authors:** Shek Man Chim, Kristen Howell, Alexandros Kokkosis, Brian Zambrowicz, Katia Karalis, Elias Pavlopoulos

## Abstract

The limited translatability of preclinical experimental findings to human patients remains an obstacle for successful treatment of brain diseases. Relevant models to elucidate mechanisms behind brain pathogenesis, such as human cell-based systems, and support successful targeting and prediction of drug responses in humans are in urgent need given the species differences in brain and blood-brain barrier (BBB) functions. Here, we examined and advanced a Brain-Chip which recapitulates aspects of the human cortical parenchyma and the BBB in one model. We utilized human primary astrocytes and pericytes, hiPSC-derived cortical neurons, hiPSC-derived brain microvascular endothelial-like cells, and included for the first time on chip hiPSC-derived microglia. Using TNFα to emulate neuroinflammation, we demonstrate that our model recapitulates *in vivo*-relevant responses. Importantly, we show microglia-derived responses, highlighting the ability of the model to capture cell-specific contributions in human disease associated pathology. We next tested BBB crossing of human transferrin receptor antibodies and conjugated adeno-associated viruses, and demonstrate successful *in vitro/in vivo* correlation in identifying crossing differences. These findings highlight the potential of the Brain-Chip as a time-saving and reliable model to support therapeutic development for brain diseases.

## INTRODUCTION

The aging global population and the increasing prevalence of age-related CNS diseases, such as Alzheimer’s and other neurodegenerative diseases, pose significant social and economic challenges. In the U.S.A alone, the economic burden of these diseases reached USD 305 billion in 2020 and is projected to soar to USD 1.1 trillion by 2050 [1,2].

Despite significant advancements in animal models and the valuable insights they have provided into brain physiology, data from animal research have often failed to predict human clinical trial outcomes, underlying the very limited availability of effective drugs for a number of brain diseases [3–7]. A major contributor to this limited translatability is the brain’s complexity and its function, which involves dynamic signaling between the numerous neuronal circuits and interactions of neurons with their surrounding cells, such as microglia and astrocytes, and the vasculature [3,6,8–10]. This inherent complexity of the brain along with species-specific differences in its function pose substantial challenges to the *in vivo* investigation of the contribution of the different cell types in the mechanisms underlying brain diseases and, therefore, the identification of specific druggable targets for patients. Furthermore, it hampers our understanding of the interactions of the Blood-Brain-Barrier (BBB) with the parenchymal cells and its potential role in disease pathogenesis and effective therapeutic targeting [10–13].

The BBB is a complex network of endothelial cells, pericytes and astrocytes with a number of species-specific features [14–22]. The tight junctions between the endothelial cells exert stringent control over the passage of molecules and harmful substances circulating in the bloodstream, ensuring CNS homeostasis and protection against neurotoxic insults. Therefore, much effort is devoted to the understanding of the biology of the human BBB and to developing strategies to overcome its restrictive properties for the successful delivery of brain therapeutics [13,23–25].

Considering these challenges, the low success rate in translating experimental findings into effective patient treatments is not unsurprising and underscores the urgent need for development of more effective models for testing CNS drugs before the first-in-human studies [3,26]. Addressing the challenge of modelling the BBB and the complexity of brain function, necessitates the development of models that may recapitulate critical aspects of their physiological complexity, thereby increasing the chances of successfully predicting drug responses in humans. This need is further endorsed by the recent FDA Modernization Act 2.0, which authorizes the use of certain alternatives to animal testing, including cell-based and computer models, to obtain an exemption from the FDA to investigate the safety and effectiveness of a new drug (congress.gov/bill/117th-congress/senate-bill/5002 ; [27]).

Recent advancements in human microphysiological systems (MPS) and Organ-Chip technologies are emerging as promising approaches to achieve this goal and provide robust and reproducible systems for drug development [6,7,28]. Microengineered Organ-Chips enable recreating a more physiological microenvironment, including co-culture of relevant cells on tissue-specific extracellular matrices (ECM), exposure to continuous flow and other *in vivo*-relevant mechanical forces such as fluidic shear stress [28–30]. Several BBB-Chip models have been designed to reconstitute the neurovascular interface [31–35] and have shown potential for exploring BBB permeability of compounds [36], nanoshuttles [37] or endothelial cell receptor-mediated transcytosis for putative brain shuttle application [38]. However, the majority of these models have not included combinations of all key BBB cell types with neurons and microglia, critical for the function of the neurovascular unit and better emulation of *in vivo* conditions [12,39]. In addition, omitting key brain parenchymal cell types limits the potential for characterization of the brain cell populations targeted by the BBB-crossing therapeutic, and early identification of therapeutic range and potentially adverse effects.

In this study, we aimed to develop a Brain-Chip to bridge some of these gaps. Using a commercially available platform, we developed a chip that includes key cell types of the cortical neurovascular unit in a single model and has a BBB with physiologically relevant permeability. We utilized human wild-type primary astrocytes and pericytes, human induced pluripotent cell (hiPSC)-derived cortical Glutamatergic and GABAergic neurons to better mimic the neurocircuitry, hiPSC-derived brain microvascular endothelial-like cells (iBMECs), and, for the first time reported on a Brain-Chip, hiPSC-derived microglia. We provide strong evidence that our model shows *in vivo* relevant responses to pharmacological and inflammatory challenges, including disruption of the BBB, and can identify cell-specific contributions. Most importantly, we demonstrate *in vivo* relevant specificity and sensitivity in screening hTfR1-based BBB-crossing therapeutic strategies, while enabling evaluation of target engagement in brain parenchymal cells. Altogether, our data demonstrate the value of using a human brain cell-based microphysiological system (Brain-Chip) as a reliable and time-sensitive model to support therapeutic development and provide mechanistic insights into human brain diseases, an unmet medical need.

## RESULTS

### Development and characterization of human cortical Brain-Chip model

To generate our Brain-Chip model, we first seeded the top channel (referred to as the “brain” channel) with human primary astrocytes and pericytes, human iPSC-derived cortical Glutamatergic (excitatory) and GABAergic (inhibitory) neurons to better mimic the neurocircuitry of the human brain cortex, and hiPSC-derived microglia, which is the first time reported in the system. Previous Brain-Chip and BBB-Chip models used a microglial cell line derived from human fetal brain primary microglia (CRL-3304) [40,41] or no microglia at all [34,42]. We chose to utilize hiPSC-derived microglia to develop a model that would enable future examination of the effects of CNS disease-associated genetic mutations in microglia cells and their contributions to cell-cell interactions and disease pathology, a role increasingly apparent in recent years [43–47].

Twenty-four hours after seeding the brain channel (Day 1), we seeded the bottom channel (referred to as the “vascular” channel) with iBMECs. On Day 2, the Brain-Chips were connected to the automated microfluidics system using a constant fluid flow of 30 μl/hr for both the brain and vascular channels (Supplementary Figure 1).

To confirm the attachment and presence of all cell types in the chip, we performed immunocytochemistry after six days in microfluidics, using the following cell-type-specific markers: Glucose transporter 1 (GLUT1) for iBMECs, Glial fibrillary acidic protein (GFAP) for astrocytes, Neural/glial antigen 2 (NG2) for pericytes, Ionized calcium binding adaptor molecule 1 (Iba1) and Cluster of Differentiation 68 protein (CD68) for microglia, and the neuronal dendritic marker MAP2 (microtubule-associated protein 2) for the detection of neurons. All cells were present and uniformly distributed throughout the respective channel (Figures 1A and B for iBMECs and brain cells, respectively). Using immunofluorescence labeling, confocal microscopy, and digital 3D image reconstruction, we also show that the astrocytes extend end-feet-like processes through the pores of the membrane reaching the endothelial cells in the vascular channel (Figure 1C), consistent with previous studies [34,40] and the role of astrocytes in the formation of the neurovascular unit and a tight BBB [48].

**Figure 1.**
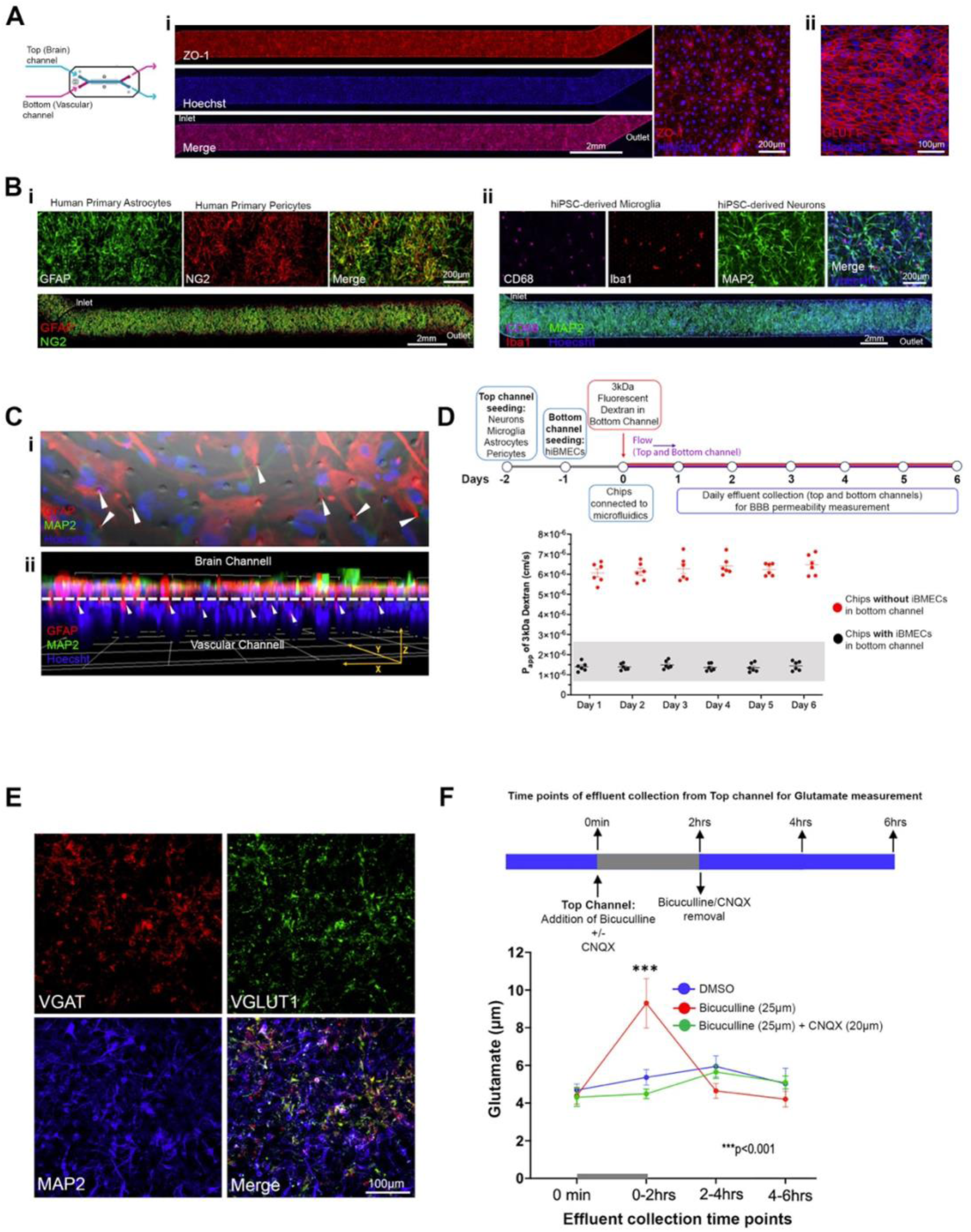
Characterization of the human cortical Brain-Chip. **(A)** Representative confocal images showing the hiPSC-derived microvascular endothelial-like cells (iBMECs) attached to the porous membrane (vascular channel). (i) Immunostaining against the tight junction marker ZO-1. Stack of Z-series of the vascular channel (left) and high magnification optical section of ZO-1 staining (right) are shown. (ii) Immunostaining against the brain microvascular endothelial cell marker GLUT1 (stack of Z-series). **(B)** Confocal images of astrocytes (GFAP) with pericytes (NG2) (i), and neurons (MAP2) with microglia (Iba1 and CD68) (ii) attached to the porous membrane in the brain channel. Confocal images (stack of Z-series) of the entire brain channel (top) and high-magnification confocal optical sections (bottom). All cell types were present and uniformly distributed along the entire brain channel. **(C)** (i): confocal micrograph (stack of z-series) showing immunofluorescence staining against GFAP (astrocytes) and MAP2 (neurons) coupled with phase contrast for visualization of the porous membrane. (ii): Digital 3D reconstruction of z-series image stacks showing the Brain-Chip from the side. The interrupted line indicates the location of the porous membrane separating the brain from the vascular channel. The nuclear staining (Hoechst) in the vascular side indicates the iBMECs. GFAP signal is detected in the vascular side (arrows). Arrows in both images indicate the astrocytic endfeet passing through the 7μm pores extending into the vascular channel. **(D)** Schematic representation of experimental design and averaged data (ii) from quantitative barrier function analysis via apparent permeability (Papp) to 3 kDa fluorescent dextran crossing through the vascular to the brain channel on days 1 through 6 in microfluidics. Chips with and without iBMECs were examined (N=6 chips/group). Each data point represents an individual chip. Graph: Mean±SEM. Shaded box: range of Papp values shown in animal models. **(E)** Confocal images showing GABAergic (VGAT) and Glutamatergic neurons (VGLUT1) in the brain channel of the chip. All stainings were performed in Brain-Chips after six days in microfluidics. **(F)** Examination of functional connectivity between GABAergic and glutamatergic neurons using pharmacology and extracellular glutamate measurements. The experimental design (top) and extracellular glutamate quantification (Mean ± SEM) for the indicated time points and treatments are shown. N=3 chips/group, ***P < 0.001, two-way ANOVA and post hoc Tukey’s test.

To ensure formation of a functional barrier, we first examined the formation of tight junctions by iBMECs using immunostaining against ZO-1 (Zona Occludens-1), a main marker of tight junctions. The iBMECs formed a uniform layer of tight junctions throughout the vascular channel (Figure 1A). Next, we measured the barrier’s apparent permeability (Papp) for a 3kDa fluorescent dextran, starting at Day 1 in microfluidics and continuing for six days (Figure 2D). Papp ranged between 1.0×10^-6^ cm/sec to 1.7×10^-6^ cm/sec, values comparable with those shown in previous studies with Brain-Chips and in rodent models (Figure 2D) [40,41,49,50].

**Figure 2.**
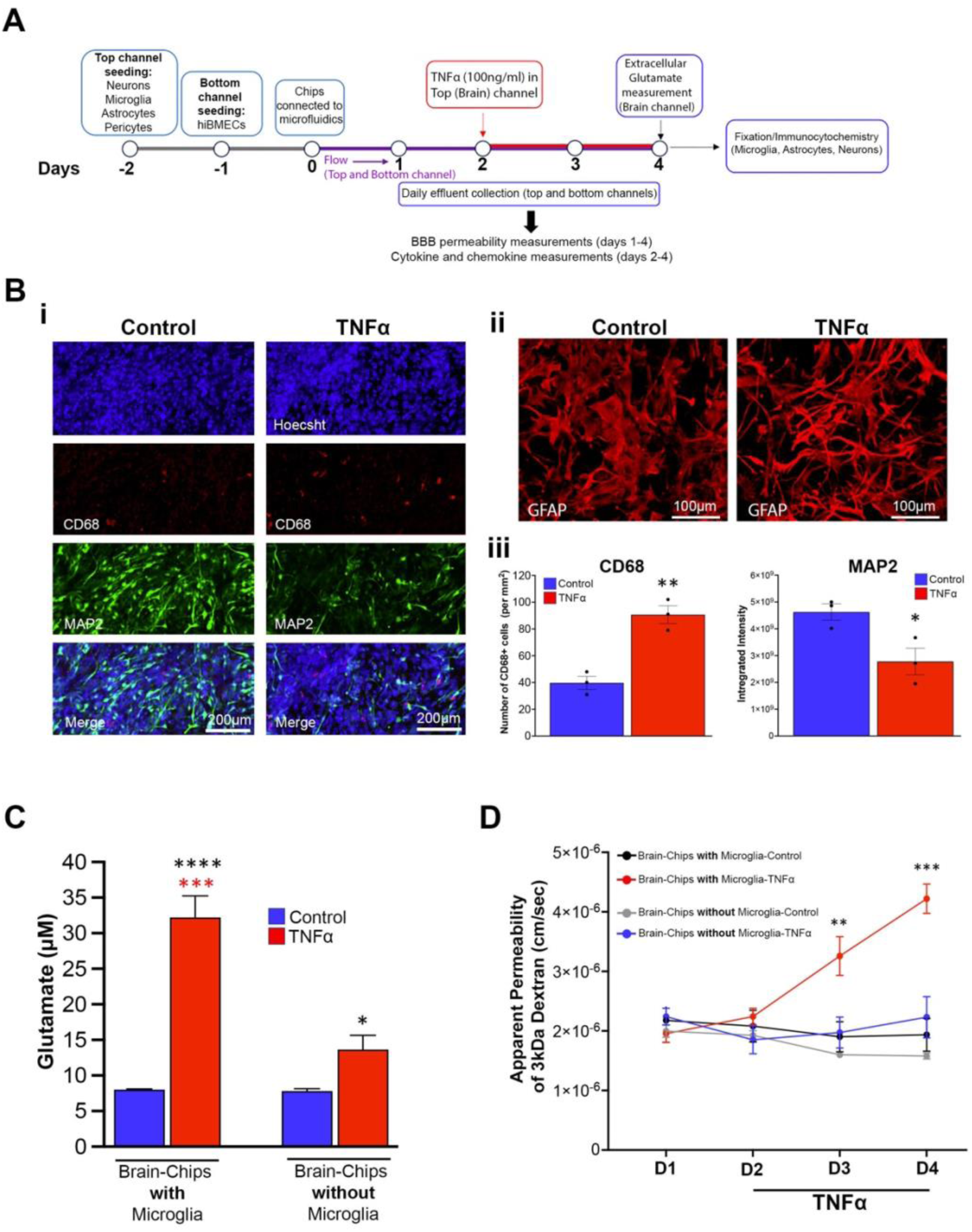
TNFα-induced neuroinflammation and BBB disruption in the Brain-Chip. **(A)** Outline of experimental design. Beginning on day 2 in microfluidics, 100ng/ml of TNFα were dosed in the brain channel and replenished 24 hours later. Chips dosed with PBS were used as control. Immunocytochemistry and extracellular glutamate measurements were performed on Day 4. Effluents were collected daily from Day 1 to Day 4 for BBB permeability assay (days 1-4) and cytokines/chemokines analysis (days 2-4). **(B)** Representative confocal images of microglia (CD68) and neurons (MAP2) (i) and astrocytes (ii). (iii) Averaged data (Mean ± SEM) of number of CD68+ cells and MAP2 intensity. TNFα treatment increases the numbers of CD68 positive cells, indicative of microglial reactivity a. The signal intensity of the neuronal dendritic marker MAP2 is decreased suggesting neuronal dysfunction. High resolution stacks of z-series from brain channel areas (50% coverage of the channel) were analyzed for each chip. N=3 chips/treatment. Confocal images of all chips used for MAP and CD68 analysis can be found in Supplementary Figure 2. The morphology of reactive astrocytes upon TNFα exposure changes from a polygonal to a more elongated state (see Supplementary Figure 3 for additional chips and images). **(C)** Averaged data (Mean ± SEM) of extracellular glutamate measurements in brain effluents collected at the end of the experiment (day 4). N=3 chips/group. Red asterisks: comparison with TNFα-treated chips without microglia. Black asterisks: comparison with respective control. **(D)** Apparent permeability (Papp) of the barrier across days (Mean ± SEM). Papp on day 2 was measured immediately before the TNFα perfusion. Chips with microglia, N=4; Chips without microglia, N=3. (B-D)*p<0.05, **p< 0.01, ***p < 0.001, p<0.0001, one-way (B and C) or two-way (D) ANOVA with post hoc Tukey’s test (significantly different compared to all other groups).

We confirmed the presence of Glutamatergic and GABAergic neurons in our chips by immunostaining against the Vesicular Glutamate Transporter 1 (VGLUT1) and the Vesicular GABA Transporter (VGAT), respectively (Figure 1E). We then tested whether these two types of neurons form functional synaptic connections and communicate. Since the chip is a closed system and electrophysiology is not possible, we used pharmacology and pulse-chase experiments. Neuronal and circuit excitability result from the synergistic action of excitatory and inhibitory inputs. Accordingly, blockade of inhibitory transmission results in enhanced excitatory transmission and increased glutamate release and extracellular glutamate levels [51–53].After four days in microfluidics, GABAergic transmission in the Brain-Chips was blocked by local (brain-channel) transfusion of the GABAA receptor blocker, bicuculline (20 μΜ). Brain channel effluents for glutamate measurement were collected immediately before the addition of bicuculline, at the end of its perfusion, which lasted 2 hours, and two and four hours later. As shown in Figure 1F, extracellular glutamate was significantly increased 2 hours after the application of bicuculline and returned to baseline levels two hours after the removal of the drug. Importantly, the effect of bicuculline on extracellular glutamate was abolished when the drug was administered together with CQNX, an AMPA receptor antagonist, in a parallel experiment (Figure 1F). This indicates that the increase in extracellular glutamate upon bicuculline was due to enhancement of excitatory transmission and glutamate release from the glutamatergic neurons.

In summary, our data demonstrate good reconstitution of our Brain-Chip model with all the seeded cortical brain parenchyma cell types present, with functional inhibitory and excitatory neuronal connections and circuitry, and a tight BBB, for at least six days in microfluidics.

### The Brain-Chip model recapitulates TNFα-induced neuroinflammation and provides evidence of microglia-specific responses

Previous studies using the Brain-Chip with the same cell types, but with a human microglial cell line, have shown that the system is functional and responsive to challenges associated with brain pathology, such as TNFα, and consequent induction of neuroinflammation, as demonstrated by assessment of three cytokines 2 days after exposure to TNFα [40]. To validate our model and, importantly, the functionality of the hiPSC-derived microglia used in it, we followed a similar approach, which is outlined in Figure 2A. Briefly, Brain-Chips were perfused with TNFα directly into the brain channel at a concentration of 100 ng/mL starting on day 2 in microfluidics for two days. The concentration of TNFα was the same as that used in previous studies and has been shown to induce the release of proinflammatory factors and compromise the integrity of the BBB [40,54–56]. Effluents for analysis were collected on day 2, just before TNFα dosing, and then every 24 hours for two more days (days 3 and 4), at which point the chips were fixed for immunofluorescence staining. Chips dosed with PBS were used as control. The brain channel effluents were collected for examination of cytokines and chemokines. Extracellular glutamate was also measured in the brain effluent collected on day 4. Compared to control chips, we observed that exposure to TNFα resulted in: 1) microglia activation, as evidenced by the increased number of cells positive to CD68, which is a lysosomal marker of microglia indicative of microglial reactivity (Figure 2B and Supplementary Figure 2), 2) transition of the morphology of astrocytes from a polygonal to a thinner and more elongated shape with longer processes, suggesting reactive astrogliosis (Figure 2B and Supplementary Figure 3), 3) reduction of the neuronal dendritic marker MAP2, indicating reduced neuronal complexity and potential neuronal dysfunction (Figure 2B and Supplementary Figure 2), 4) a significant increase in extracellular glutamate levels (Figure 2C), 5) disruption of the integrity of the BBB, as evidenced by the significant increase in its apparent permeability to 3kDa dextran (Figure 2D), and 6) induction of pro-and anti-inflammatory cytokines and chemokines (Figure 3).

**Figure 3.**
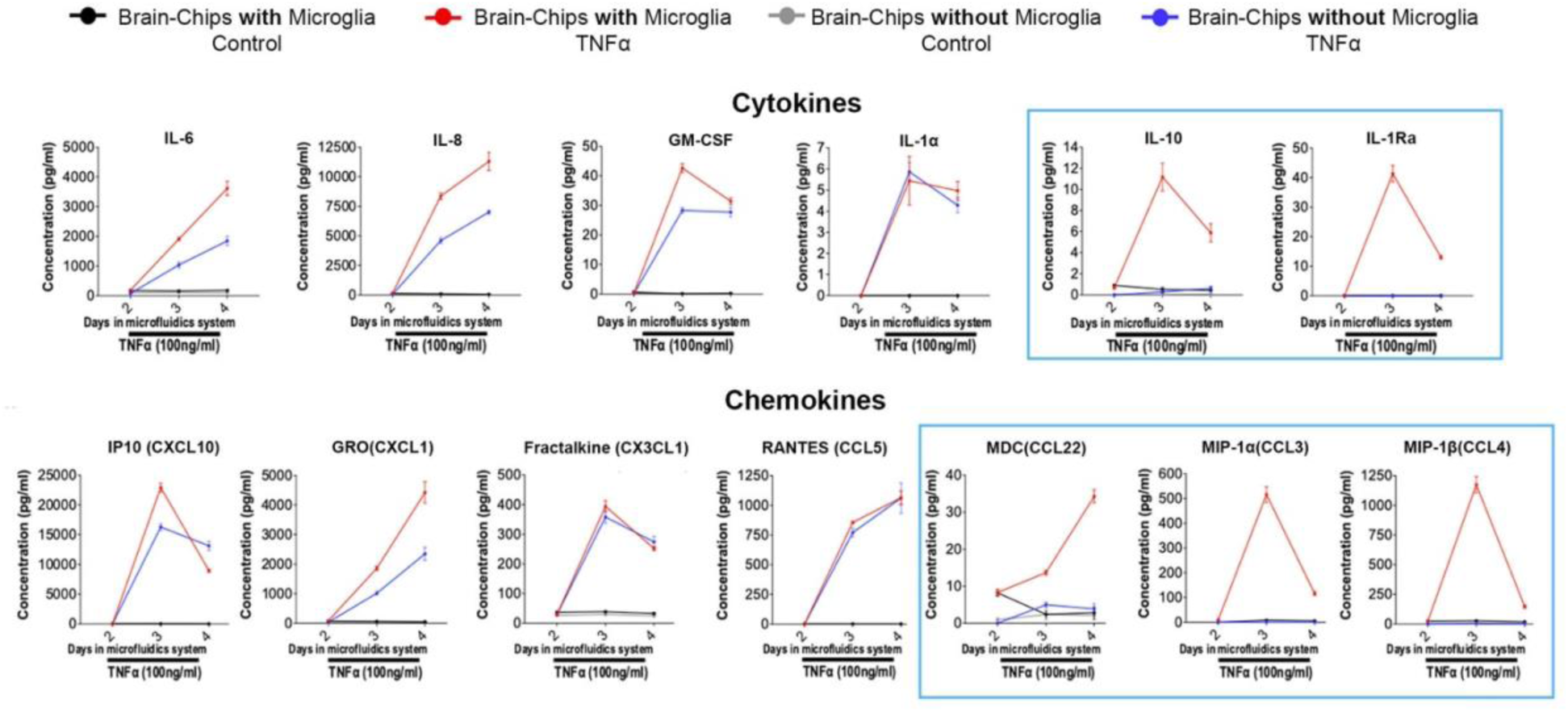
TNFα-induced secretion of cytokines and chemokines and contribution of microglia. Longitudinal analysis of cytokines and chemokines in brain channel effluents collected at the indicated time points. Effluent collection on day 2 was performed immediately prior to TNFα dosing (baseline levels of cytokines and chemokines). Brain-Chips with and without microglia were examined to determine their contribution to the observed inflammatory responses. All other cell types (astrocytes, pericytes, Glutamatergic and GABAergic neurons, brain microvascular endothelial-like cells) were present in the chips. Brain-Chip (with and without microglia) treated with PBS were used as controls. Graphs: averaged data (Mean ± SEM) from 4 chips for each group. Microglial-specific responses are indicated by the blue rectangles.

All the effects of TNFα observed in our Brain-Chip are consistent with published *in vitro* and *in vivo* studies and, therefore, demonstrate that our model is functional and capable of emulating pathological conditions affecting brain function [40,57–71]. Of note, the daily collection of effluents enabled identification of the temporal profile of cytokine and chemokine responses to TNFα (Figure 3).

In addition to examining Brain-Chips with all the cell types included in the model, we also examined, in parallel, Brain-Chips without microglia. We found that the TNFα-induced increase in extracellular glutamate was ameliorated and the disruption of the BBB was abolished in Brain-Chips without microglia (Figure 2C and D). This is consistent with the important contribution of these cells to neuroinflammation and BBB disruption and in agreement with published works [12,61,70,72–75]. Additionally, we mapped the contribution of microglia to the temporal profile of cytokine and chemokine changes induced by TNFα. Indeed, a subset of cytokines (IL-6, IL-8, GM-CSF) and chemokines (GRO) continued to be induced by TNFα, but these responses were significantly lower in the chips without microglia. Notably, some cytokines and chemokines were not affected by the lack of microglia (IL-1a, IP-10, Fractalkine, and RANTES), whereas the TNFα-mediated induction of the proinflammatory IL-10 and anti-inflammatory IL-1Ra and the chemokines MDC, MIP-1α, and MIP-1β was completely abolished (Figure 3). The microglial-specific induction of IL-10, IL-1Ra, MDC, MIP-1α, and MIP-1β levels are compatible with their expression profile, which is restricted to microglia and macrophages [76], in further support of the specificity of the responses obtained with the Brain Chip.

In summary, our validation data demonstrate the functionality of our Brain-Chip and the hiPSC-derived microglia used in the system for the first time, and the capability of our model to emulate pathological conditions affecting the brain. Moreover, we demonstrate that our model has the resolution to determine cell-specific contributions, for better understanding the dynamic cell-cell interactions in brain pathogenesis.

### Human TfR1 but not mouse TfR1-specific antibody BBB crossing in the Brain-Chip

A promising method extensively studied to overcome the BBB challenge for brain-targeting therapeutics is crossing the BBB via endothelial receptor-mediated transcytosis. Thus, we examined whether our cortical human Brain-Chip could serve as a screening platform for testing BBB-crossing therapeutic strategies, focusing on the human transferrin receptor 1 (hTfR1), a widely used approach for targeted brain drug delivery [77].

First, we examined the specificity of our model to detect hTfR1-mediated BBB crossing. After reconstitution and 24-hour incubation in microfluidics (day 1), chips were perfused in the vascular channel for 24 hours with human antibody specific for human TfR1 [78] or mouse TfR1 [79] (10μg/ml concentration in the endothelial cell culture media). Chips perfused with human IgG1 (isotype) or treated with PBS were used as negative controls. A 3kDa fluorescent dextran was perfused in the vascular channel since the beginning of incubation in microfluidics (Day 0) for daily BBB permeability measurements to ensure that all chips had a tight BBB with comparable permeability (Days 1 and 2; Figure 4B).

**Figure 4.**
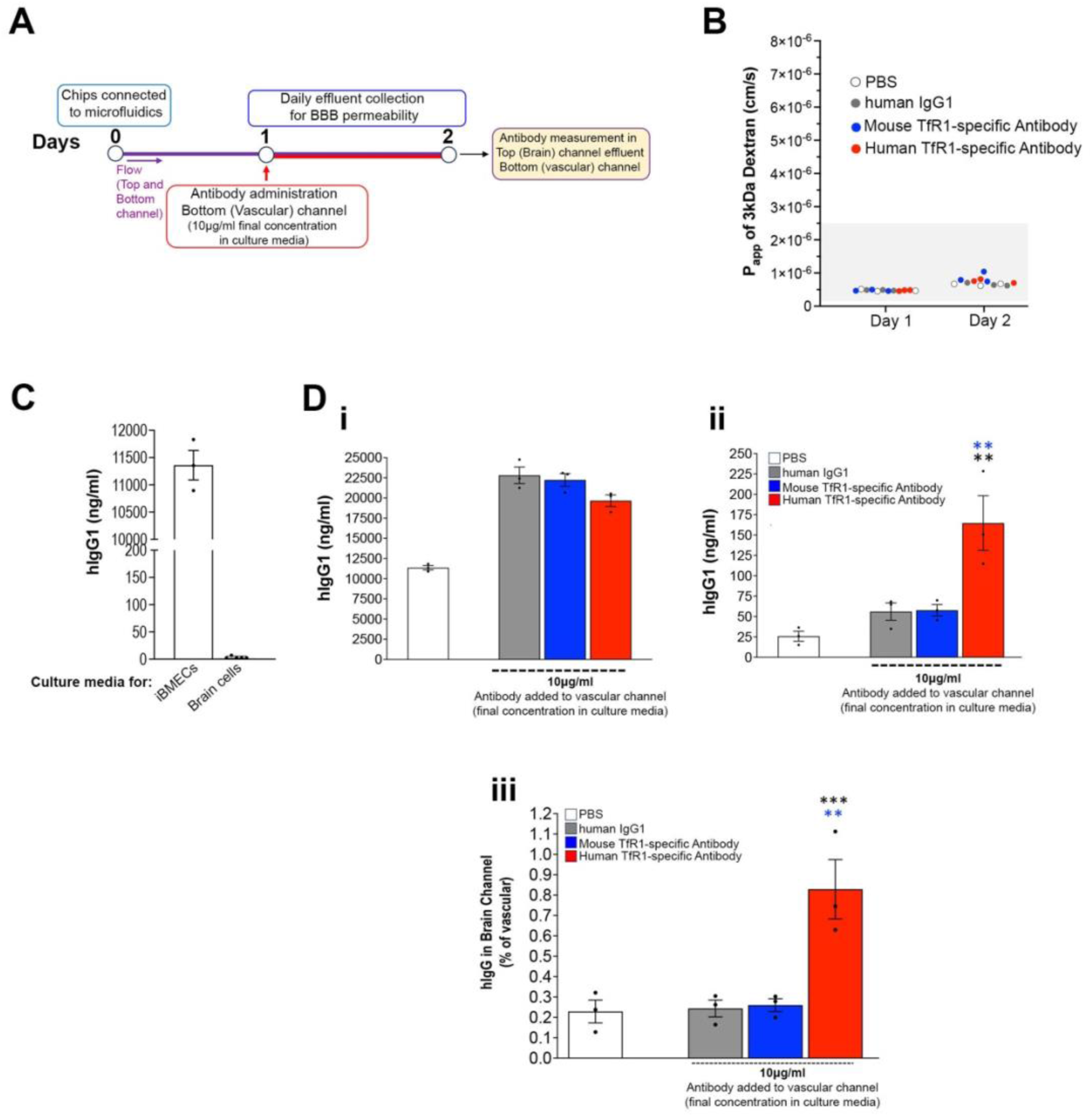
Specificity of the Brain-Chip for human Transferrin Receptor-mediated BBB crossing. **(A)** Schematic representation of experimental design. On Day 1 in microfluidics, the vascular channel was perfused with human antibody specific for human TfR1 or mouse TfR1, or isotype control antibody (human IgG1) at a final concentration ∼10 μg/ml. PBS was used as a baseline control. On Day 2, effluents from the brain channel were collected for hIgG1 measurements. The permeability of the barrier was examined on days 1 and 2 by measuring its Papp to 3kDa fluorescent dextran perfused to the vascular channel since the beginning of incubation in microfluidics. **(B)** All chips had a tight barrier with similar Papp values, which were within the range of those shown in published works and in rodent models (Shaded box) . Each data point represents an individual chip. **(C)** Measurement of hIgG1 levels in culture media of vascular and brain channels prior to the perfusion of the TfR1 antibodies. hIgG1 was detected in the vascular maintenance medium, as it contained 2% human serum. Data points represent three individual measurements. **(D)** Quantification of hIgG1 in the vascular channel media prior to perfusion (i) and in brain channel effluents collected 24 hours post-dosing. The percentage of vascular hIgG1 detected in the brain channel for each treatment group was also calculated (iii). Averaged data (Mean ± SEM) and individual chip values are shown in (i-iii). N=3 chips per group. **p < 0.01, ***p < 0.001, one-way ANOVA with post hoc Tukey’s test. Blue asterisks: comparison with mouse TfR1. Black asterisks: comparison with hIgG1.

The vascular channel media initially contained detectable human IgG1 (∼11.3 ± 0.27 μg/ml), due to the supplementation of 2% human serum. After antibody dosing, hIgG1 levels increased in the media accordingly and were similar across the treatment groups (Mean + SEM; μg/ml for hIgG1, mTfR1 and hTfR1: 22.4 ± 1 μg/ml, 22.9 ± 0.74 μg/ml and 20.2 ± 0.7 μg/ml, respectively; Figures 4C and D). The brain channel media lacked human serum and had undetectable hIgG1 before dosing.

Twenty-four hours post-treatment, we collected brain channel effluents and measured hIgG1. Chips dosed with the hTfR1 antibody had significantly more hIgG1 in the brain channel compared to chips dosed with the mTfR1 antibody (Mean ± SEM; μg/ml; 150.8 ± 0.03 and 57.6 ± 0.007; hTfR1 and mTfR1, respectively). hIgG1 levels in the brain channel of mTfR1-treated chips were similar to those in chips with isotype control antibody (64.8 ± 0.01 μg/ml) and slightly higher than PBS-treated groups (25.9 ± 0.006 μg/ml) (Figure 4D). Normalized data showed that the percentage of vascular hIgG1 detected in the brain channel of the PBS-treated chips, which is likely due to paracellular transport, was indistinguishable from mTfR1 and isotype control treated chips (Mean ± SEM; 0.24 ± 0.05 %, 0.26 ± 0.04% and 0.28 ± 0.05%; PBS, hIgG1 and mTfR1, respectively). In contrast, this percentage was significantly greater than all other groups in chips treated with hTfR1 antibody (0.756 ± 0.1%). This effect was not due to changes in BBB integrity, as we identified no differences in permeability between chips (Figure 4B). These data demonstrate human TfR1-specific BBB crossing in the human cortical Brain-Chip model.

### The Brain-Chip identifies BBB crossing differences between hTfR1 antibodies

After establishing that the Brain-Chip can detect hTfR1-specific BBB crossing, we examined its ability to discern varying levels of hTfR1 antibody permeation. For this purpose, we tested internally generated hTfR1-specific antibodies. These antibodies had previously been evaluated in human TfR1 knock-in (KI) mice as hTfR1:AAV9GFP formats (conjugated to AAV9 expressing GFP). Their BBB crossing ability was determined based on immunohistochemical detection of GFP in the mouse brain parenchyma and blinded scoring of the GFP signal (Supplementary Figure 4).

For our Brain-Chip experiment, we used purified antibodies and selected a range of them based on their BBB crossing properties. Specifically, we tested the best crosser from the *in vivo* screening (REGN1), a good and a moderate crosser (REGN5 and REGN12, respectively), and a non-crosser (REGN28). The experimental design was similar to that used for testing the specificity of the Brain-Chip with two modifications. First, we replaced the culture media of the iBMECs with serum-free media in order to eliminate the serum-derived IgG levels and, therefore, precisely measure the dosed antibodies. Second, we perfused the antibodies for eight hours (Figure 5A). We have found this time point to give similar results with the 24 hour-perfusion (data not shown). The chips were equilibrated for 24 hours in microfluidics prior to perfusion with the antibodies, which were dosed in the vascular channel at a concentration of 10 μg/ml in the culture media. Brain channel effluents were collected at the end of the antibody perfusion. As shown (Figure 5B), REGN1 antibody levels in the brain channel effluent were nearly double of those of the REGN5 antibody, which were significantly higher than REGN12 (Mean ± SEM; ng/ml; REGN1, REGN5 and REGN12, respectively: 656.4 ± 20.7, 345.3 ± 40.6 and 184.1 ± 25.4). REGN28, a non-crosser *in vivo*, was detected at low levels (50.5 ± 11.1 ng/ml) comparable to the isotype control antibody (59.6 ± 10.1 ng/ml), suggesting paracellular transport. Normalization with vascular channel antibody levels showed similar results (Figure 5B; Mean ± SEM; Percentage of dosed antibodies detected in the brain channel; REGN1, 5.6 ± 0.23; REGN5, 3.0 ± 0.35; REGN12, 2.2 ± 0.26; REGN28, 0.35 ± 0.07; isotype control, 0.5 ± 0.10). Our data align with those in hTfR1 KI mice, demonstrating that the human Brain-Chip model can accurately detect different levels of BBB permeation of hTfR1 antibodies and predict *in vivo* responses. Importantly, this prediction was possible in just 4 days, including the two days required for the seeding of the cells and the preincubation of the chips under flow for one day, a significantly faster process compared to animal studies.

**Figure 5.**
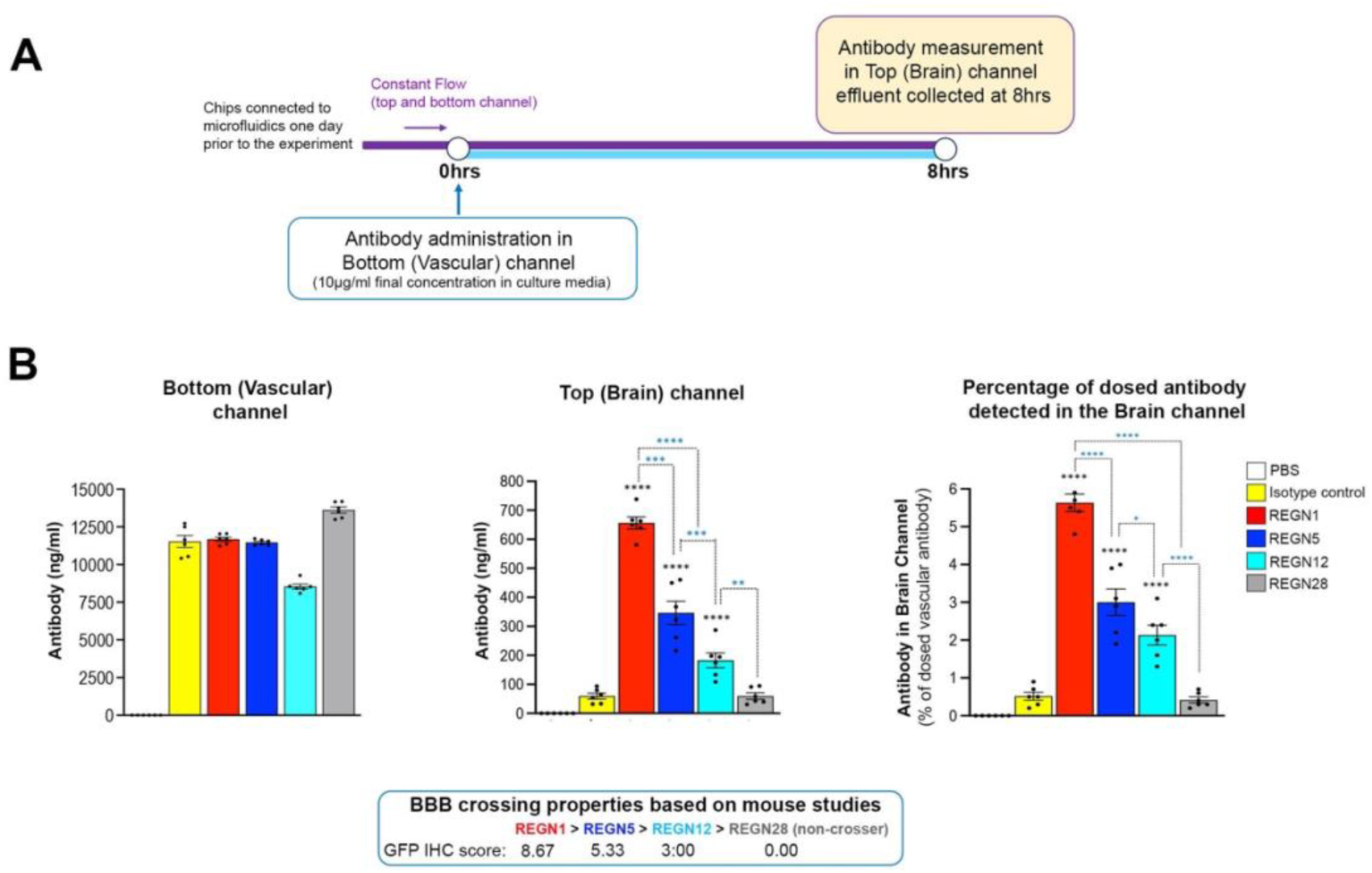
Human Brain-Chip has the resolution to identify BBB crossing differences between human TfR1 antibodies. **(A)** Outline of experimental design. The chips were connected to microfluidics for one day prior to antibody administration of human TfR1 antibodies with varying BBB crossing properties *in vivo*. The vascular channel was dosed with the antibodies in serum-free culture media at 10μg/ml. PBS and isotype antibody were used as negative controls. After 8 hours, effluents from the brain channel were collected for antibody measurements. **(B)** Antibody quantification data. Mean ± SEM and individual chip values are shown. Antibody quantification in vascular channel media prior to perfusion (left) and in the brain channel effluent 8 hours post-dosing (middle) are shown. Right graph shows the percentage of the dosed antibodies detected in the brain channel. Four human TfR1 antibody clones with different BBB crossing ability were examined (REGN1, REGN5, REGN12 and REGN28). The clones had been tested in mice as conjugates to AAV9 expressing GFP and their BBB crossing ability was determined based on immunohistochemical detection and quantification of the GFP signal in the brain parenchyma. The numeric immunohistochemical scores and their order based on their BBB crossing ability are shown below the graphs. N=6 chips/group. *p < 0.05, **p < 0.01, ***p< 0.001, ****p < 0.0001, one-way ANOVA with post hoc Tukey’s test. Black asterisks: comparison with isotype antibody control. No differences in BBB permeability were observed between groups (Supplementary Figure 5A).

### The human Brain-Chip detects BBB crossing differences between hTfR1 antibody-conjugated AAV9

To further validate the Brain-Chip, we re-examined the internally developed hTfR1 antibodies, now conjugated to AAV9 expressing GFP, the same format used in mice. To refine our model’s resolution, we included an additional antibody, REGN25, which exhibited low BBB crossing in mice [BBB crossing ability *in vivo*: REGN1 > REGN5 > REGN12 > REGN25 > REGN28 (non-crosser)]. Chips treated with PBS, AAV9 without any conjugated antibody, and AAV9 conjugated with an antibody specific to the liver-expressed Asialoglycoprotein receptor 1 (hASGR1) served as controls.

Twenty-four hours post-calibration in the microfluidics (Day 1), the chips were exposed to the virus in the vascular channel for two consecutive days (6.94×10^6^ viral genomes/μl; 1×10^10^ viral genomes in total). On Day 3, the vascular media were replaced with fresh media without virus, and the chips were incubated for two additional days (Day 3 to Day 5) to allow maximal GFP expression. Endothelial and brain parenchymal cells were then collected and GFP protein levels measured using automated ELISA (Figure 6A and Materials and Methods). Total protein in cell lysates was comparable between the groups (Supplementary Figure 5B).

**Figure 6.**
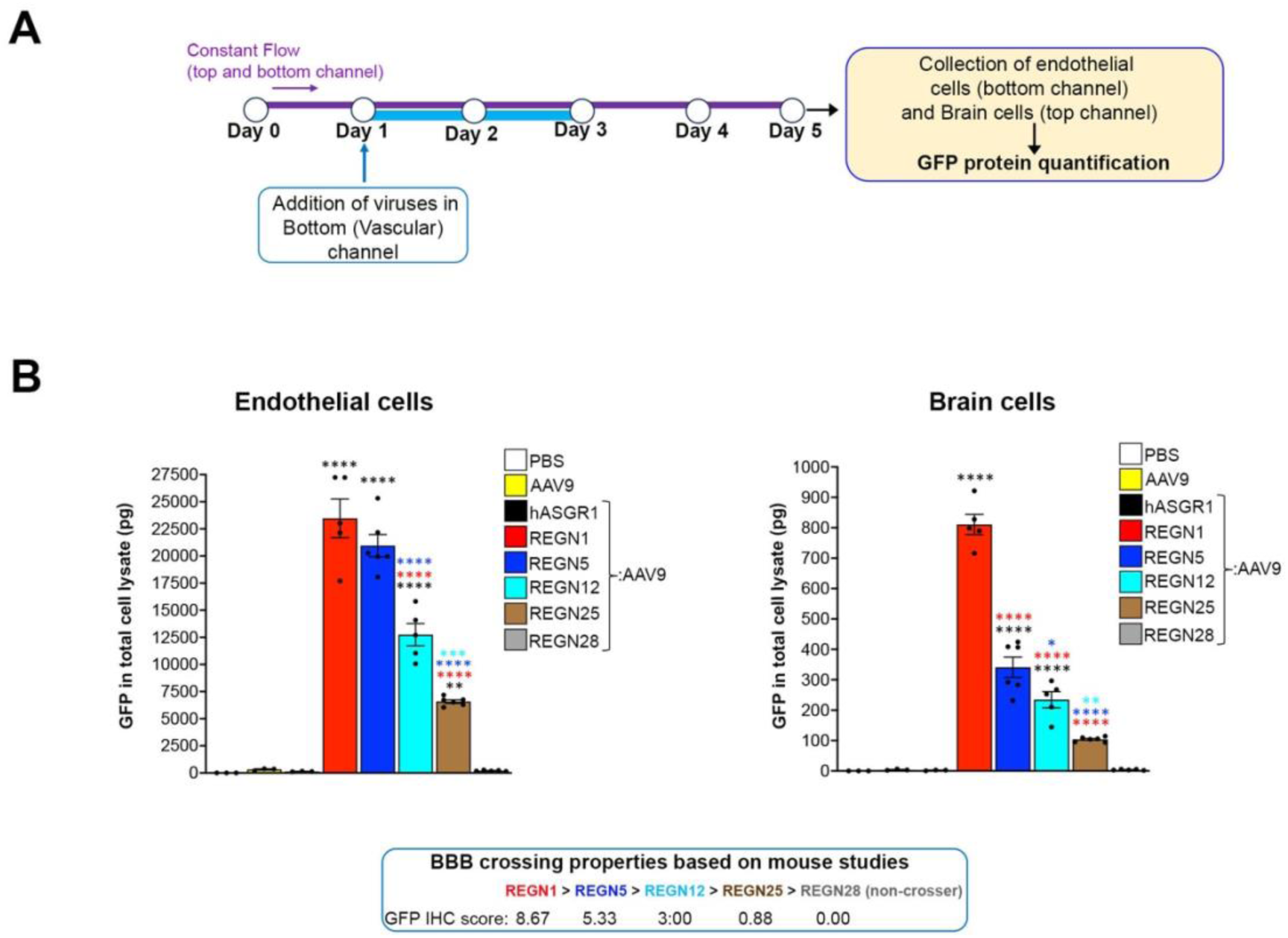
The human Brain-Chip detects different levels of permeation hTfR1 antibody-conjugated AAV9. **(A)** Schematic of experimental design. AAV9 expressing GFP and decorated with hTfR1 antibodies were perfused in the vascular channel for two days (Day 1-3). On day 5, the endothelial-like cells from the vascular channel and the brain channel cells were collected for GFP protein analysis. **(B)** Graphs showing averaged data (Mean ± SEM) and individual chip values of GFP protein quantification analysis in endothelial and brain cell lysates, as indicated. The hTfR1 antibody-conjugated viruses had been tested *in vivo* for their BBB crossing abilities, based on GFP signal intensity scoring in the mouse brain parenchyma, as shown below the graphs. We examined AAV9 conjugated with the same hTfR1 clones tested as purified antibodies in the chip (REGN1, REGN5, REGN12, REGN28) plus viruses conjugated with the antibody clone REGN25, which exhibited low BBB crossing *in vivo*. Unconjugated AAV9 and hASGR1-conjugated viruses were used as negative controls. hTfR1 antibody-conjugated AAV9: N=6 chips for each group; PBS, AAV9 and hASGR1-AAV9: N=3; *p < 0.05, **p < 0.01, ***p < 0.001, ****p < 0.0001, one-way ANOVA with post hoc Tukey’s test. Black asterisks: comparison with PBS, AAV9 and hASGR1-AAV9. Red asterisks: comparison with REGN1. Blue asterisks: comparison with REGN5. Light blue asterisks: comparison with REGN12. All chips had a tight barrier with comparable Papp (Supplementary Figure 5A). The amount of total protein in endothelial and brain channel cell lysates was comparable between groups (Supplementary Figure 5B)

As shown in Figure 6B, the total GFP in endothelial cells of chips treated with REGN1:AAV9 and REGN5:AAV9 was comparable, indicating similar virus uptake (Mean ± SEM; 23.02 ± 1.78 ng and 20.10 ± 1.02 ng, respectively). The GFP in endothelial cells from chips treated with REGN12:AAV9 (12.69 ± 1.03 ng) was significantly lower compared to REGN1:AAV9 and REGN5:AAV9, but significantly higher compared to REGN25:AAV9 (6.61 ± 0.16 ng). GFP levels in endothelial cells from chips treated with REGN28:AAV9 (non-crosser), hASGR1:AAV9, and unconjugated AAV9 were minimal (Mean (ng) ± SEM; 0.18 ± 0.021, 0.13 ± 0.013, 0.35 ± 0.055, respectively).

In the brain parenchymal cells, GFP level difference closely matched the *in vivo* BBB crossing studies and our Brain-Chip experiment with purified antibodies. REGN1 showed the greatest GFP expression, followed by REGN5, REGN12, and REGN25. The non-crosser REGN28 and all control viruses showed no GFP expression [Mean (pg) ± SEM; 798.3 ± 33.29, 349.4 ± 33.51; 252.7 ± 26.45, 105.1 ± 3.15, respectively]. GFP expression differences were statistically significant. A parallel experiment where chips were perfused with the same viruses and cells collected at the end of viral dosing (Day 3) for viral genome measurements showed consistent results with the GFP expression analysis and BBB crossing properties in mice (Supplementary Figure 5B).

In summary, our data demonstrate the successful development of a human cortical Brain-Chip model with strong potential for modelling disease-associated phenotypes for mechanistic studies and as a screening platform for hTfR1-based therapeutic strategies

## DISCUSSION

Enhancing the likelihood for successful translation of preclinical findings into clinical settings necessitates an in-depth understanding of the mechanisms driving disease development and progress in patients, along with capabilities for the best possible prediction of safety and efficacy of therapeutics before the first-in-human studies. Addressing such requirements poses a particular challenge for brain diseases and cannot be easily met using animal models due to limited predictive and face validity given the remaining unknowns in disease pathogenesis and related phenotypes. Key reasons include species differences in brain and BBB biology, as well as the impact of genetics including the frequent lack of similarities in genotype/phenotype associations between humans and experimental animal models. Along the same lines, several disease-risk variants lie in non-coding regions, which are markedly different between humans and animals [80]. Potential conformational and tissue expression differences of disease-associated targets can also limit the effectiveness of animal models in predicting successful therapeutic targeting [3,6,81,82].

Human cell-based systems recapitulating the human brain microenvironment are emerging as a promising approach to elucidate brain disease mechanisms and enable the establishment of reliable and time-saving drug screening platforms [28,83,84]. Given the BBB’s importance and the critical roles of neurons, glial, immune, and vascular cells in brain physiology in health and disease through both cell-autonomous and non-cell autonomous effects [44,85–94], significant efforts have been directed toward developing platforms incorporating all key cell types and vasculature in a single model [28,95,96] emulating critical elements of the organ microenvironment.

In the present study, we utilized a commercially available Organ-Chip platform to develop a human cortical Brain-Chip that includes five major brain cell types and recreates the neurovascular unit and the BBB to further expand on the capabilities of previously reported *ex vivo* systems. Cortical neurons, astrocytes, microglia, and pericytes form the brain parenchymal side (top channel of the chip), while brain microvascular endothelial-like cells make up the vascular side (bottom channel). Astrocytes extend end-feet-like processes to the vascular side through membrane pores, indicating the recreation of the abluminal aspect of brain microvessels. The astrocytes and microglia remain in a quiescent state, suggested by the very low or nearly undetectable baseline levels of secreted cytokines and chemokines (Figure 3). Over six days of culture under constant flow, the endothelial-like monolayer maintains a tight barrier, comparable to animal models [49,50].

Besides combining key brain parenchymal cell types and the BBB in a single model, we demonstrated functional connectivity between inhibitory (Glutamatergic) and excitatory (GABAergic) cortical neurons by pulse-chase experiments with bicuculline and CNQX and extracellular glutamate measurements. The baseline levels of glutamate across all chips and experiments were comparable (6-7 μM; Figures 2 and 3) and within the range observed in the human brain cortex (0.1-20μM, depending on the cortical area) [97,98], indicating a physiologically relevant circuitry and excitatory/inhibitory (E/I) balance. Upon TNFα challenge, extracellular glutamate levels rose to concentrations typically used for the induction of excitotoxicity in neuronal cultures [99–101], consistent with the role of TNFα in inducing glutamate-mediated excitotoxicity [66,67] and the disturbed E/I balance observed in several brain diseases and neuroinflammation [102–104]. Thus, our Brain-Chip enables measurements of a neurocircuitry-related and physiologically-relevant endpoint for mechanistic and pharmacological studies. Our results with bicuculline demonstrate the chip’s suitability for pulse-chase experiments and its sensitivity to capture responses to transient pharmacological challenges. The return of the extracellular glutamate levels to baseline after the wash-out of bicuculline indicates healthy and plastic neurons and neural circuitry. It also proves that the setup of this engineered model permits well-controlled and precise pharmacological manipulations without disturbing the cells, most likely due to the constant and automated control of flow, an advantage over Transwells, organoids and other static culture systems.

To validate the *in vivo* relevant responses of our Brain-Chip to disease-associated challenges, we assessed responses to TNFα exposure through the brain channel. We confirmed glutamate excitotoxicity, microglial and astrocytic reactivity, activation of the secretion of cytokines and chemokines, neuronal damage, and compromised BBB, consistent with previous studies and the role of TNFα in neuroinflammation, a major component in the pathogenesis of CNS diseases [105–112]. Importantly, we demonstrate microglia-specific inflammatory responses and microglia-dependent BBB disruption. This aligns with published studies [73] and further supports the critical role of microglia in the neuroimmune mechanisms involved in human brain pathology [44]. Our data also demonstrate the advantage of our model to map cell-specific contributions and dynamic cell-cell interactions critical in disease-associated pathology. Additionally, we were able to generate a temporal secretory profile for cytokines and chemokines over several days, before and during TNFα exposure. Such fine mapping of inflammatory responses is important for disease modelling and comparison of genetic variants associated with brain disease traits. The ability of the chip to capture time-dependent phenotypic and functional changes within the same sample chip (or same “individual”), in response to experimental interventions is another of its main advantages compared to static multicellular systems.

Preclinical testing of brain-targeting therapeutics, particularly biologics, remains challenging. Given the limitations of animal studies, developing clinically relevant human *in vitro* models for predicting penetration, efficacy, and investigating transcytosis mechanisms for BBB-crossing therapeutic strategies and drug development is of major need. Brain endothelial cell receptors and receptor-mediated transcytosis (RMT) represent an emerging strategy for BBB crossing and potentiating therapeutic delivery to the brain [23,113,114]. Strategies utilizing RMT involve creating a complex between the therapeutic modality and a receptor-targeting entity, such as an endogenous receptor ligand, mimetic peptide ligand, or receptor-targeting antibody. The therapeutic can be chemically linked to the entity (i.e. chemical compound or siRNA linked to a receptor-specific antibody) or incorporated in a vehicle decorated with the RMT-targeting entity, such as in the case of adeno-associated viruses (AAVs) expressing a therapeutic protein [113,115–118]. Human Transferrin receptor 1 (hTfR1) is among the most studied RMT targets in brain endothelial cells [77,116,119–121]. Mice have provided important insights into the therapeutic potential of the approach, although they are not hTfR1-specific. hTfR1 KI mice have been generated to overcome this issue and are currently used in proof-of-concept studies for therapeutic testing. This important animal model has great potential, but the question of its predictive value due to differences in the biology of the BBB between rodents and humans remains open. Considering that the human TfR1 acts in the context of a mouse cell environment, BBB crossing studies in the hTfR1 KI mouse would be greatly benefited by complementary studies in exclusively human cell-based systems. *In vivo* screening of hTfR1-specific entities and therapeutic modalities can be also challenging. Effective BBB crossing and brain-targeting is a multistep process including efficient uptake by endothelial cells, efficient transcytosis, and successful release to the brain parenchymal side and engagement in the targeted cells. Meeting these criteria requires screening numerous versions of the examined entity; for example, multiple antibody clones and formats in the case of hTfR1-targeting antibodies. Such *in vivo* screening efforts take a lot of time (several months), are labor-intensive, and, importantly, require a very large number of mice, which is not aligned with the 3R principles of animal use in biomedical research – reduction, refinement, replacement – aiming for significant reduction of the use of animal models in preclinical studies [122].

We examined BBB crossing differences of hTfR1 antibodies in two formats: 1) conjugated to AAV9, the format used for *in vivo* testing, and 2) unconjugated. This was possible because of the versatility of the chip, which enables collection of cells and effluents from the brain and vascular sides for appropriate readout measurement methods for each antibody format. The comparable results from the two experiments demonstrate the robustness, reproducibility and sensitivity of the model. In addition, the selected readouts, which are quantitative, are more reliable than qualitative methods such as measuring changes in immunostaining intensity. Furthermore, our results with hTfR1:AAV9 illustrate the Brain-Chip’s suitability for analyzing endothelial cell uptake and determining target engagement in the brain parenchymal compartment. With the right design and bioassays, future studies that combine biochemical analysis with cell imaging will allow for the examination of the cell types targeted by the tested therapeutic modalities as well as the prediction of target engagement, safety and, potential efficacy in one single study. Considering the time required to generate and test multiple antibody clones and formats, and the large number of animals necessary for such experiments, our human Brain-Chip holds great potential to facilitate screening efforts and advance therapeutic development. Furthermore, comparison between basal (healthy) and disease states, such as in neuroinflammation or genetic models, might provide disease-specific insights that could further support drug development.

The Brain-Chip we describe in this study, shows the sensitivity and specificity to qualify as a screening platform for hTfR1-based BBB crossing therapeutic strategies. We demonstrated that the model could identify BBB crossing differences between different hTfR1 antibodies tested in hTfR1 KI mice, aligning with the associated *in vivo* findings. By measuring BBB permeability during these experiments, we ensured a tight barrier with comparable permeability in all examined chips, indicating that the observed differences were due to transcytosis rather than a leaky barrier.

Our data demonstrate several advantages of the Brain-Chip. However, the model in its current stage has certain limitations. It is composed of five different cell types from different human donors, and a mix of primary (astrocytes and pericytes) and hiPSC-derived cells (neurons, microglia and iBMECs). This set up is less than ideal for disease modelling studies. Although there has been significant progress in developing protocols for differentiating human iPSCs into microglia, astrocytes, pericytes and oligodendrocyte-lineage cells, the functionality of each of these cells in complex *in vitro* models requires experiments beyond the scope of this study. As a first step, we selected to use hiPSC-derived microglia and examine their functionality in the chip, which is the first time. The cells are functional and exhibit neuroinflammatory responses upon exposure to TNFα, consistent with *in vivo* data and a recent study using a similar Brain-Chip configuration, but with the human microglial cell line CRL-3304 [40]. Our Brain-Chip established a tight barrier with *in vivo* relevant permeability. However, the exact identity of iBMECs is still debatable [123], so we chose to refer to them as endothelial-like cells. The iBMECs used in our study express genes regulating key BBB attributes, such as genes encoding for the tight junction proteins ZO-1 (*TJP1*), Occludin (*OCLN*) and Claudin (*CLDN*), solute carrier transporters, ATP-binding cassette transporters and GLUT1 (*SLC2A1)*, and the TfR1-encoding gene, *TFRC* [124] and data not shown).

The closed format of the Organ-Chip design does not support the examination of Transendothelial Electrical Resistance (TEER), an *in vitro* measure of electrical resistance across a cell layer considered a sensitive measure of barrier integrity [125,126]. Therefore, we measured the Papp of the barrier using fluorescent dextran, an accepted alternative used in several published works using Organ-Chips and animal models [34,40,41,49,50,127]. Papp is a relevant surrogate to TEER and a corollary in the clinic, enabling *in vivo* comparison of barrier integrity, which is not possible with TEER measurements.

Despite these caveats, our Brain-Chip model in its current configuration provides unique advancements compared to previous Brain-Chip and BBB-Chip models. It confirms functionality at the pharmacological level as shown both for BBB crossing and neuronal responses to relevant pharmacologic challenges. Further, it recapitulates phenotypic changes and responses to neuroinflammation-relevant challenges in a time-dependent manner, while highlighting microglia-dependent cytokine release, a major area of investigation for a great number of brain diseases. These findings provide evidence that the human Brain-Chip holds strong potential for modelling and therapeutic development for neuroinflammatory and neurodegenerative diseases. Importantly, our Brain-Chip enables timesaving, specific, and accurate detection of different levels of permeation of *in vivo*-tested hTfR1-specific antibodies and associated therapeutic modalities (AAV9), demonstrating its value in facilitating and advancing the development of human brain-targeting therapies. Future advancements should incorporate oligodendrocyte precursor cells (OPCs) and oligodendrocytes (OLs) and develop Brain-Chip models with all cell types fully isogenic and derived from hiPSCs. This would enable detailed mapping of the mechanism of action of disease-associated genetic variants, including determination of cell-autonomous and non-cell autonomous effects, to further inform cell-specific targeting, and identification of biomarkers to guide patient stratification. The continuous improvements in cell differentiation protocols, the scientific community’s and pharmaceutical companies’ growing interest in human cell systems [28,128] and the FDA Modernization Act provide realistic optimism for the engagement of human cell systems in facilitating drug development and fighting brain diseases.

## MATERIALS AND METHODS

### Cell culture

Commercial human iPSC-derived cortical Glutamatergic and GABAergic neurons, and human primary astrocytes were purchased from NeuCyte (SynFire® Co-Culture kit; Cat.# 1010-7.5). The cells were cultured in Neucyte cell maintenance media. Primary human brain pericytes (isolated from human cerebral cortex tissue) were purchased from Cell Systems (Cat.# ACBRI 498) and cultured according to the manufacturer’s protocol. Human iPSC-derived microglia were purchased from Fujifilm Cellular Dynamics (Cat.# C1110).

### Differentiation of hiPSCs into human brain microvascular endothelial-like cells (iBMECs)

Human-induced pluripotent stem cells (hiPSCs) were purchased from iXCell Biotechnologies (Cat.# 30HU-002; Lot.# 400221) and maintained in mTeSR™ Plus medium (Stemcell Technologies, Cat#. 100-0276) on six-well culture plates coated with Matrigel (Corning, Cat.# 354277). Directed differentiation of hiPSCs was adapted from a previous publication (Qian et al., 2017). Briefly, hiPSCs were expanded in mTeSR™ Plus medium. The cells were then treated with 6 µM CHIR99021 (Reprocell, Cat.# 04-0004-10) in DeSR1 medium, which is composed of DMEM/Ham’s F12 (Thermo Fisher Scientific, Cat.# 11039021), 1% MEM-NEAA (Thermo Fisher Scientific, Cat.# 11140050), and 0.1 mM 2-Mercaptoethanol (Sigma, Cat.# M3148). Twenty-four hours later, the medium was replaced by DeSR2 medium [DeSR1 plus 1x B27 (Thermo Fisher Scientific, Cat.# 17504044)]; the medium was refreshed every day for a period of another 5 days. On day 6, the medium was switched to hECSR1 [Human Endothelial SFM (Thermo Fisher Scientific, Cat.# 11111044) supplemented with 20 ng/mL bFGF (R&D Systems, Cat.# 233-FB), 10 μM retinoic acid (Sigma, Cat.# R2625) and 1x B27]. On day 8, the medium was replenished with freshly prepared hECSR1. On day 9 the cells were switched to hESCR2 medium (Human Endothelial SFM supplemented with 1x B27). On day 10, the cells were dissociated with Accutase (Stemcell Technologies, Cat.# 07920) and replated in a T-75 flask coated with 400μg/mL Collagen IV (Sigma, Cat.# C5533), 100μg/mL Fibronectin (Corning, Cat.# 356008) and 20μg/mL Laminin (Thermo Fisher Scientific, Cat.# 23017015). After 20 minutes, the unattached cells were removed using Human Endothelial SFM supplemented with 5% human serum (Sigma, Cat.# P2918) and 10 µM Y27632. The iBMECs were maintained in Human Endothelial SFM supplemented with 5% human serum until seeding into the Brain-Chip.

### Brain-Chip microfabrication and Zoë^®^ culture module

Organ-Chips (Chip-S1^®^, Emulate, Inc. Boston, MA, USA) were used to setup the human Brain-Chip [40,41]. The chip is made of transparent, flexible polydimethylsiloxane (PDMS), an elastomeric polymer, and contains two parallel microchannels: the top (Brain) channel and the bottom (Vascular) channel (Dimensions: 1 x 1 mm and 1 x 0.2 mm, respectively). The two channels are separated by a thin (50 μm) porous membrane (pore diameter: 7 μm; pore spacing: 40 μm) made of PDMS (Huh et al., 2013). The co-culture area of two parallel channels is 17.1 mm^2^. Flow can be introduced to each channel independently to continuously provide essential nutrients to the cells, while effluent containing any secretion/waste components from cells is collected on the outlet of each channel separately. The Zoë^®^ culture module (Emulate) is the instrumentation designed to automate the maintenance of the chips (12 chips per module) in a controlled manner.

### Human Brain-Chip and cell seeding

Prior to cell seeding, chips were functionalized using Emulate’s proprietary protocols and reagents. Briefly, ER-1 (Emulate, Cat.# ER-105) and ER-2 (Emulate, Cat.# ER-225) were mixed at a concentration of 1 mg/mL before their addition to the top and bottom channels of the chip. The platform was then irradiated with high-power UV light having peak wavelength of 365 nm and intensity of 100 μJ/cm^2^ for 20 min using a UV oven (CL-1000 Ultraviolet Crosslinker AnalytiK-Jena: 95-0228-01). After surface functionalization, both channels of the chip were coated with Collagen IV (400 μg/mL), Fibronectin (100 μg/mL), and Laminin (20 μg/mL), and incubated at 4°C overnight. Before cell seeding, the chips were incubated at 37°C for 1 hour, followed by a PBS wash. The top (brain) channel was then seeded with a mix of hiPSC-derived glutamatergic and GABAergic neurons (4 x 10^6^ cells/ml and 2 x 10^6^ cells/ml respectively), human primary astrocytes (2 x 10^6^ cells/ml), human iPSC-derived microglia (0.8 x 10^5^ cells/ml), and primary pericytes (2 x 10^5^ cells/ml). The cells were mixed in “seeding medium” (NeuCyte) and incubated in the channel for three hours (37°C and 5% CO2). The top channel was then washed with seeding medium to remove unattached cells and incubated overnight at the same conditions. The next day, the iBMECs were seeded in the vascular channel at a density of 16 x 10^6^ cells/ml using iBMEC seeding medium (human serum-free endothelial cell medium supplemented with 5% human serum from platelet-poor human plasma (Sigma, Cat.# P2918) and allowed to attach to the membrane overnight (the bottom side of the chip was facing up during this period). The iBMECs were incubated in the channel for 3 hours, then washed with iBMEC seeding medium to remove unattached cells. After overnight incubation (37°C and 5% CO2), the chips were flipped back to their original orientation and connected to the Zoë® Culture Module. The medium supplying the brain channel was switched to maintenance medium (Neucyte), whereas the serum of the vascular medium was lowered to 2% (vascular maintenance medium). Chips were maintained under constant one-way flow (30 μl/h for both brain and vascular channel) at 37°C with 5% CO2.

### Immunofluorescence microscopy

Chips were fixed with 4% paraformaldehyde in PBS for 15 min and then washed with PBS six times. Incubation of the chips in blocking/permeabilization buffer (10% donkey serum, 0.1% Saponin in PBS) for 2 hours was the next step followed by overnight incubation at 4°C with primary antibodies diluted in blocking/permeabilization buffer (for antibody dilutions see the “Antibodies” section of “Material and Methods”). After six washes with washing buffer (PBS with 0.1% Saponin), cells were incubated for 1hr at room temperature with secondary antibodies conjugated with Alexa Fluorophores in blocking/permeabilization. The cells were washed six times with PBS. Incubation for 2 minutes with Hoechst 33342 (NucBlue™ Live ReadyProbes™ Reagent; Invitrogen, Cat.# R37605) for nuclear stain was the next step, followed by six washes with PBS and final mounting with (Ibidi, Cat.# 50001). Images were acquired with the LMS880 Zeiss confocal microscope.

### Antibodies

Chicken anti-MAP2 (Abcam, Cat.# ab5392, 1:200), mouse anti-GFAP (Abcam, Cat.# ab279290, 1:300), rabbit anti-IBA1 (Fujifilm Cellular Dynamics, Cat.# 019-19741, 1:50), mouse anti-CD68 (Abcam, Cat.# ab955, 1:100), rabbit anti-NG2 (Abcam, Cat.# ab83178, 1:100), mouse anti-VGLUT1 (Invitrogen, Cat.# MA5-31373, 1:100), rabbit anti-VGAT (Millipore, Cat.# AB5062P, 1:100), rabbit anti-GLUT1 (Abcam, Cat.# ab115730, 1:200) and rabbit anti-ZO1 (Invitrogen, Cat.# 40-2200, 1:50). Secondary antibodies conjugated with Alexa Fluor-488 (Abcam; Cat.# ab63507 and ab150073), Alexa Fluor-568 (Abcam; Cat.# ab175470) and Alexa Fluor-647 (Abcam; Cat.# ab150075 and ab150107) were used at 1:300 dilution.

### BBB permeability assay

To evaluate the integrity of the BBB, 3 kDa Dextran-Cascade Blue (Invitrogen, Cat.# D7132), was added to the vascular compartment of the Brain-Chip at 0.1 mg/ml. The effluents from both channels were collected daily for 1 or more days (depending on the experiment) for fluorescence measurements by SpectraMax M4 (Molecular Devices). The apparent permeability (Papp) was calculated based on a standard curve and using the following formula, as previously described [41].

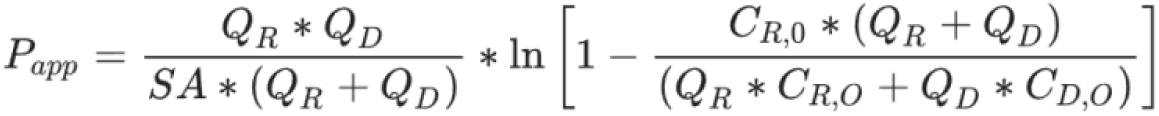

SA is the surface area of sections of the channels that overlap (0.17cm^2^), QR and QD are the fluid flow rates in the dosing and receiving channels respectively, in units of cm^3^/s, CR,O and CD,O are the recovered concentrations in the dosing and receiving channels respectively, in any consistent units.

### Pharmacological studies with bicuculline

Bicuculline to block GABAA receptors and GABAergic transmission was purchased from Torcis and used at 20 μM (Cat.# 0130). The AMPA receptor antagonist CNQX to simultaneously block glutamatergic transmission was used at 25 μM (Torcis, Cat.# 0190). Prior to the addition of the drugs, the effluents of the chips were removed from the outlets to ensure the collection of fresh effluents during the experiment. After two hours and immediately before the drug administrations, the brain channel effluents were collected for the measurement of the baseline levels of glutamate. Brain channel effluents were again collected at the end of the 2-hour perfusion of the drugs. Perfusion of the chips continued with fresh and drug-free maintenance culture media for four more hours . Brain channel effluents were collected 2 and 4 hours after the removal of the drugs. Glutamate was measured using an assay kit purchased from Abcam, (Cat.# ab83389). Chips treated with DMSO were used as controls, given that DMSO was used to reconstitute bicuculline and CNQX.

### TNF-α induced neuroinflammation

Brain-Chips were incubated in microfluidics prior to exposure to TNFα (R&D Systems, Cat.# 210-TA), which was perfused to the brain channel at 100 ng/ml concentration starting on day 2. On day 3, the brain channel media were replenished with freshly prepared TNFα. Effluents from the brain channel collected immediately before the administration of TNFα (day 2) were used for baseline measurements. During the two days of the administration of TNFα, effluents were collected on a daily basis (days 3 and 4) for analysis. Extracellular glutamate was measured in effluents collected at the end of TNFα administration (day 4) using the same kit used for the glutamate measurements in the bicuculline experiment (Abcam, Cat.# ab83389). Chips were then fixed for immunostaining and confocal microscopy. BBB permeability assays to examine the integrity of the barrier were performed on a daily basis, as described above. The levels of secreted cytokines and chemokines in the collected effluents were measured using the MILLIPLEX MAP Human Cytokine/Chemokine Magnetic Bead Panel-Immunology Multiplex Assay from Millipore (Cat.# HCYTOMAG-60K), following the manufacturer’s directions. Chips with similar culture conditions but perfused with PBS were used as controls. The experimenter was blinded to the treatment groups.

### BBB crossing studies

We tested the specificity of the Brain-Chip using a human TfR1 antibody (JCR Pharmaceuticals clone 3) [78] and a mouse TfR1 antibody (8D3 clone) [79]. The antibodies were added in vascular maintenance medium at 10 μg/ml concentration and flowed to the vascular channel at 30 μl/h for 24 hours, at which point the brain channel effluents were collected for analysis. The detection and quantification of antibodies were performed by a human IgG1-specific ELISA kit (Abcam Cat.# ab100548).

The purified hTfR1 antibodies used to test the sensitivity of the Brain-Chip to detect BBB crossing differences were internally generated (Regeneron Pharmaceuticals, Inc., Tarrytown, NY, USA). We replaced the culture media of the iBMECs with serum-free Human Endothelial SFM one hour prior to the perfusion of the antibodies. The antibodies were added in the serum-free media (10 μg/ml) and flowed to the vascular channel at 30 μl/h for eight hours at which point the brain channel effluents were collected for detection and quantification of the antibodies using ELISA (Abcam, Cat.# ab157709).

All AAV9 were obtained from the Viral Production Core at Regeneron Pharmaceuticals, Inc. (Tarrytown, NY, USA). The packaged genome contained a single-stranded GFP reporter driven by the scCBh promoter. The viruses (1×10^10^ viral genomes per chip) were added in the vascular maintenance medium and flowed to the vascular channel at 30 μL/h for 2 days. Cell lysates were collected from top and bottom channels two days after the perfusion of the viruses. GFP protein quantification was performed using the Ella™ Automated ELISA platform (Bio-techne, Cat.# SPCKB-OT-002820). For the quantification of viral genomes, DNA was collected from the cell lysates at the end of the viral perfusion using the DNA Extract All Reagents Kit from Thermo Fisher Scientific (Cat.# 4402616). Digital PCR was performed to quantify the AAV ITR2 sequence (5’-3’; Forward primer: GGAACCCCTAGTGATGGAGTT-3’, Reverse primer: CGGCCTCAG-TGAGCGA, Probe Sequence: CACTCCCTCTCTGCGCGCTCG) using the QIAcuity Probe PCR Kit (Qiagen, Cat.# 250103) and the QIAcuity Digital PCR System (Qiagen, Cat.# 911050), following the manufacturer’s protocol. The experimenter was blinded to the treatment groups.

### Statistical analysis

Statistical analysis was performed using GraphPad Prism software, applying one-way or two-way ANOVA, followed by post hoc Tukey’s tests. Statistical significance reported when p<0.05. Sample sizes are described in each figure legend.

## Data Availability Statement

The data presented in this study are available in this article.

### Acknowledgments

We are grateful to Kyle Plasterer for providing the AAVs described in our manuscript. We also thank John Dugan, Nicole Keating, and Pascaline Aimé-Wilson for their work on the mouse BBB-crossing studies and for sharing their data. Additionally, we thank Henderson Jones and Monique Nichols for quantifying the viral genomes. We extend our thanks to the Therapeutic Proteins group for producing the hTfR1 antibodies, and we are particularly indebted to Heather Bender for coordinating the antibody production and providing the antibodies.

## SUPPLEMENTARY FIGURES

**Supplementary Figure 1.**
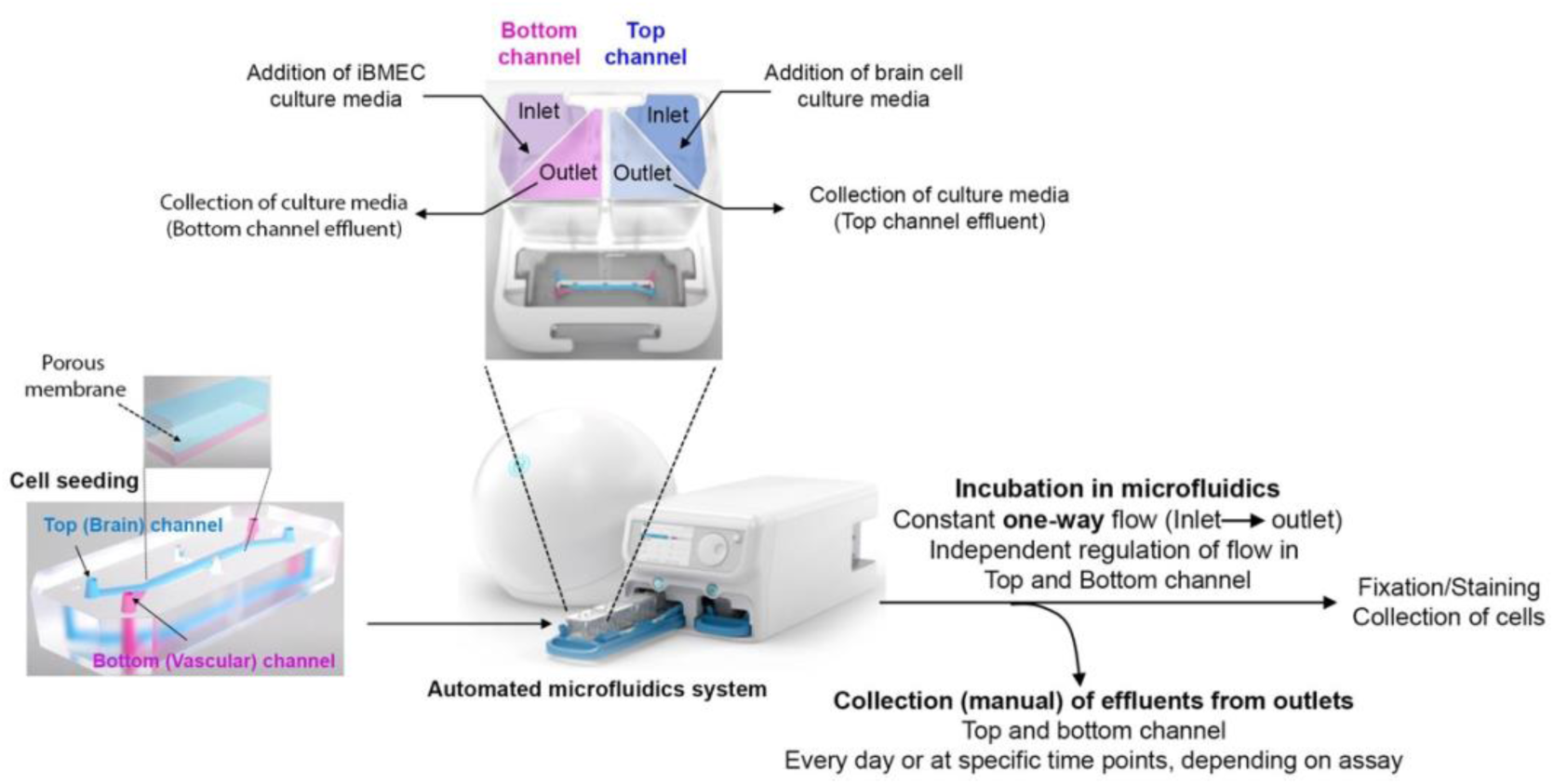
Overview of human Brain-Chip experimental setup. Endothelial cells are incorporated in the bottom (vascular) channel and are separated from the parenchymal compartment (top or brain channel) by a semi-permeable membrane coated with tissue-specific extracellular matrix. After cell seeding, the Chips are connected to the Pods which supply culture media to the top and bottom separately. Fresh culture media are added to the inlet and flow into the channels. Effluents are collected in the outlets. The chips are cultured in the automated microfluidics system which regulates a constant one-way flow in top and bottom channels independently. Effluents are collected at different time points for analysis based on the assays. At the end of each experiment, cells are fixed or collected for downstream analysis.

**Supplementary Figure 2.**
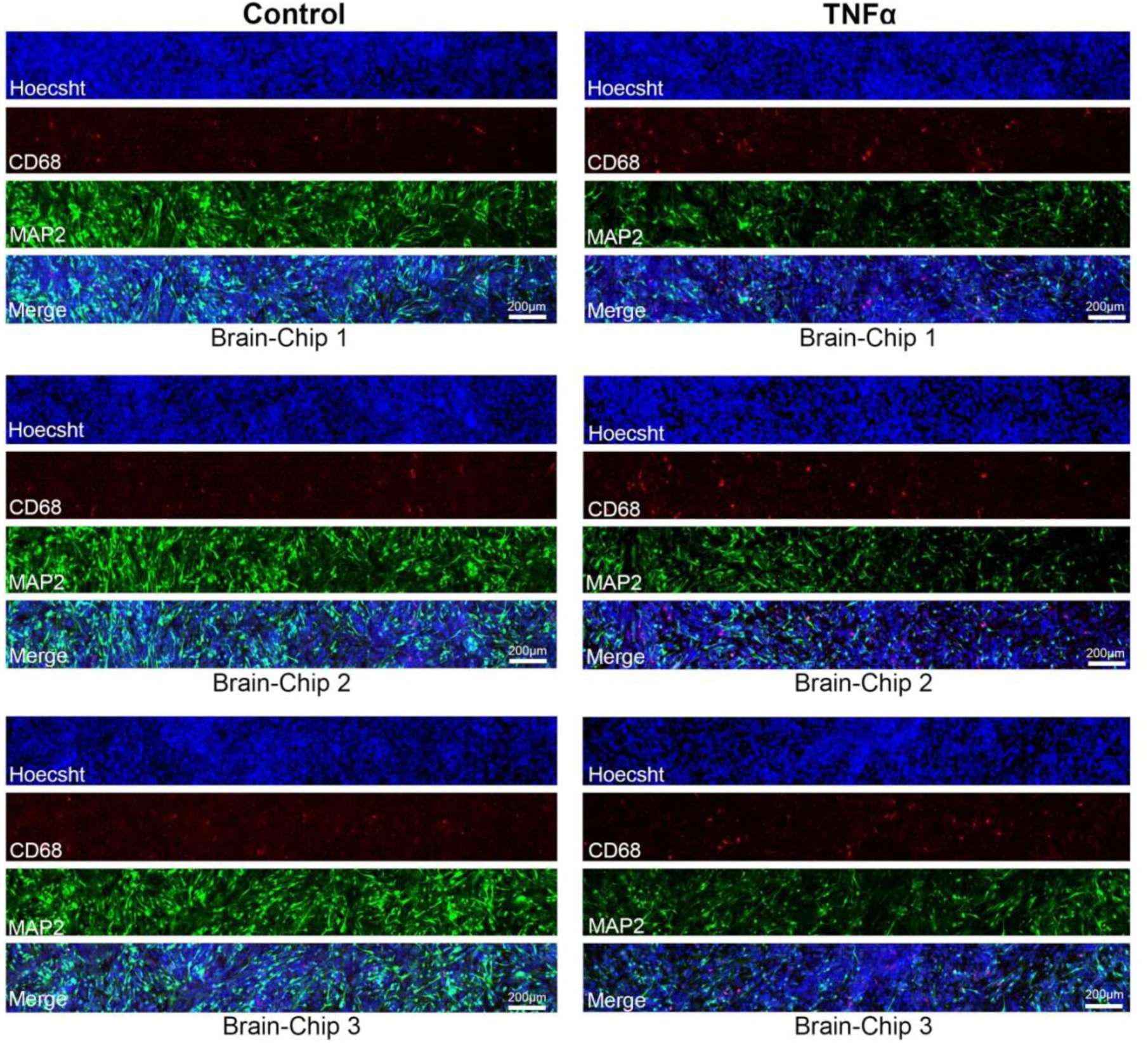
Activation of microglia and neuronal damage in the Brain-Chip upon TNFα treatment. Confocal images (stack of z-series) of microglia (CD68) and neurons (MAP2). Increase of CD68 positive cells and reduction of MAP2 indicate microglial reactivity and neuronal damage in response to TNFα, respectively. Supplementary data for Figure 2B.

**Supplementary Figure 3.**
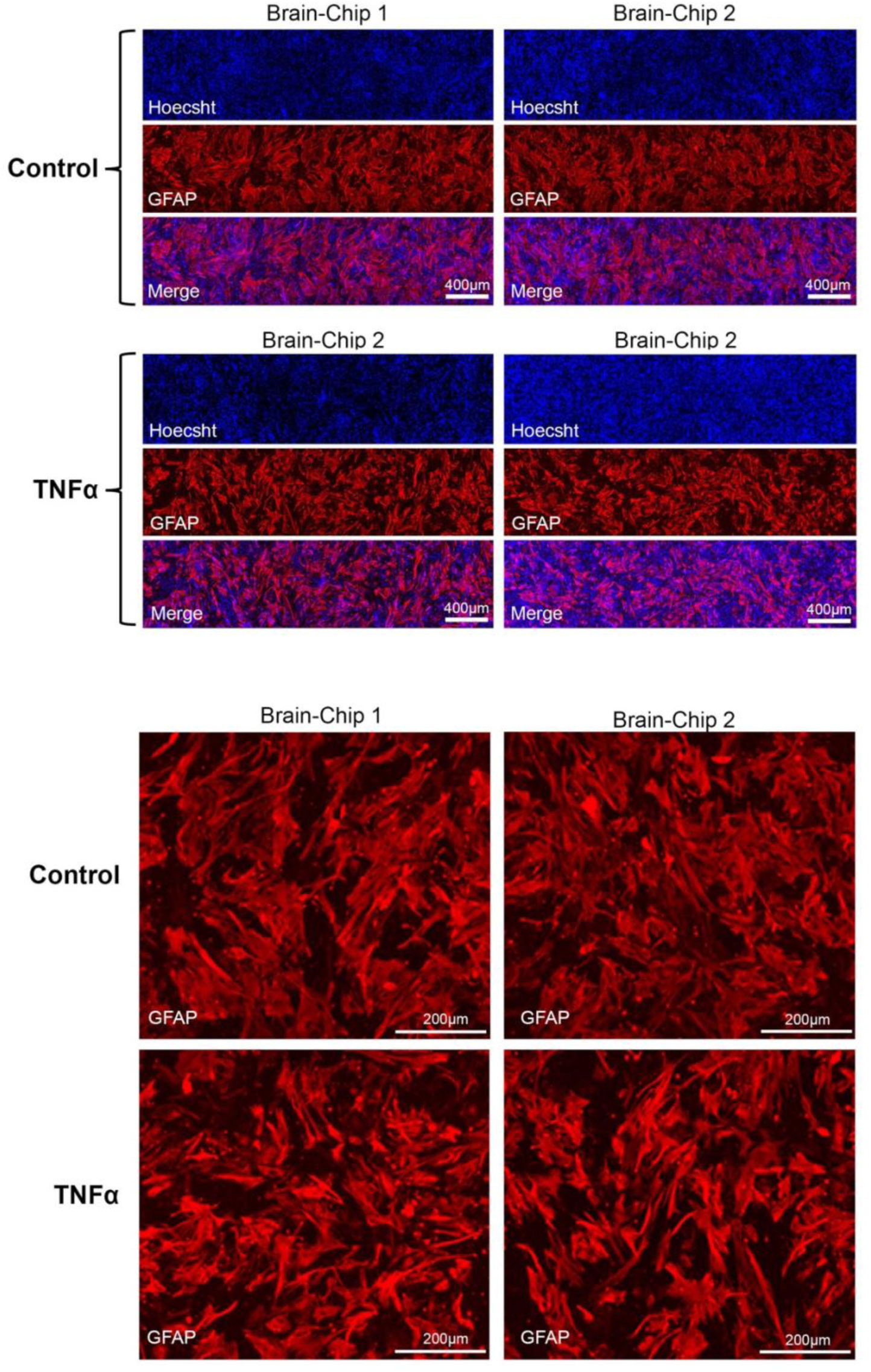
TNFα-induced morphological changes of astrocytes in the Brain-Chip. Confocal images (stack of z-series) of immunofluorescent staining against the astrocytic marker GFAP. Two chips per group were examined. Astrocytes transition from a polygonal to a more elongated shape upon exposure to TNFα. Supplementary data for Figure 2B.

**Supplementary Figure 4.**
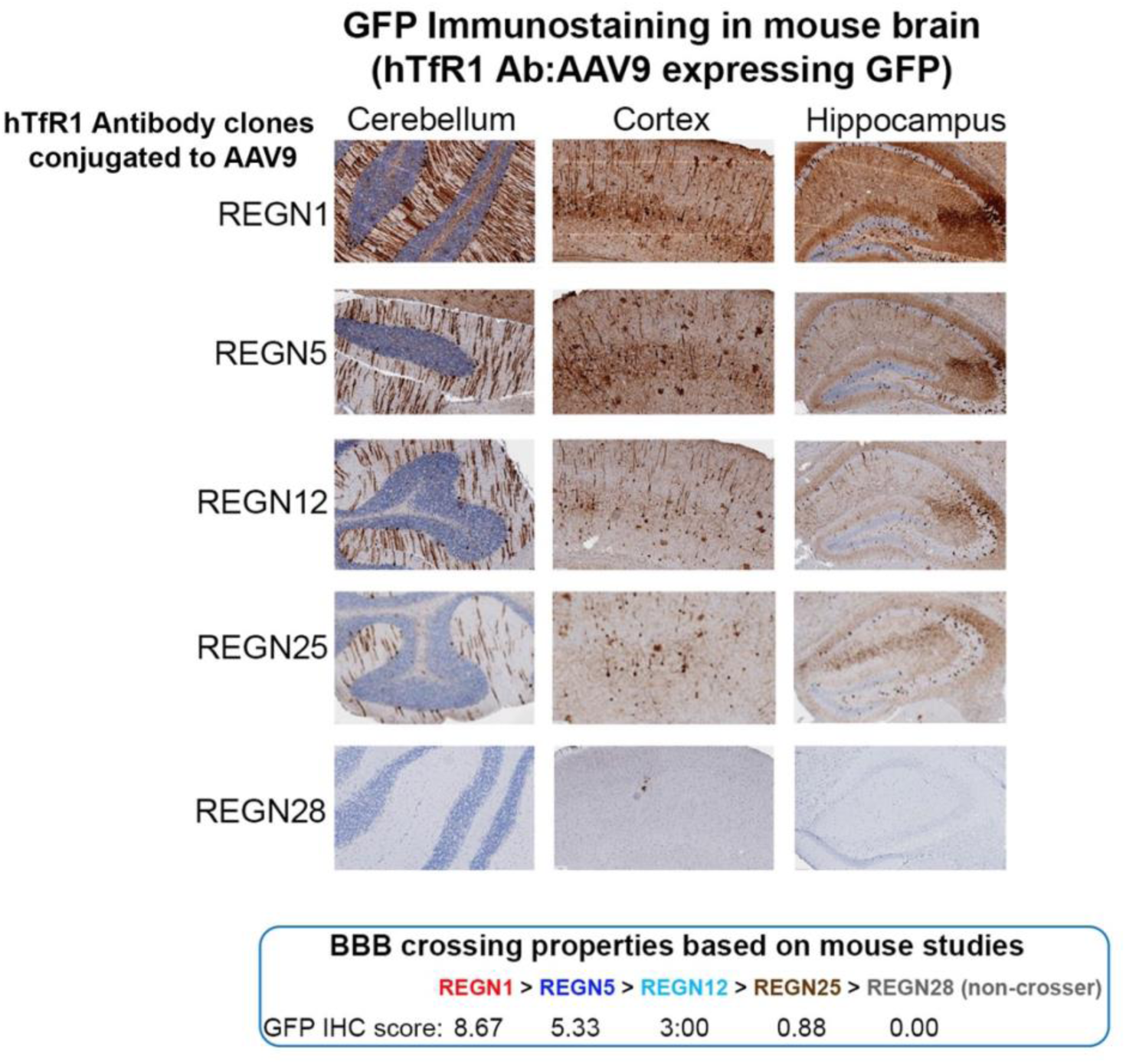
*In vivo* tested hTFR1 antibody clones examined in the Brain-Chip for BBB crossing. Representative images of GFP immunohistochemistry in different brain areas of hTfR1 knock-in mice injected i.v. with AAV9 conjugated with the indicated hTfR1 antibody clones. The crossing properties of the clones were determined based on the GFP signal in the mouse brain parenchyma and blinded quantification analysis. Score “zero” indicates no brain GFP signal and, therefore, no BBB crossing. The hTfR1antibody clones shown in the figure are part of a large *iv vivo* screening assay with thirty-two antibody clones. REGN1 was the best BBB crosser and REGN28 was among the clones that did not cross the BBB (no GFP signal). The GFP signal scoring for REGN1 and the rest of the antibody clones examined in the Brain-Chip is shown. Supplementary data for Figures 5 and 6.

**Supplementary Figure 5.**
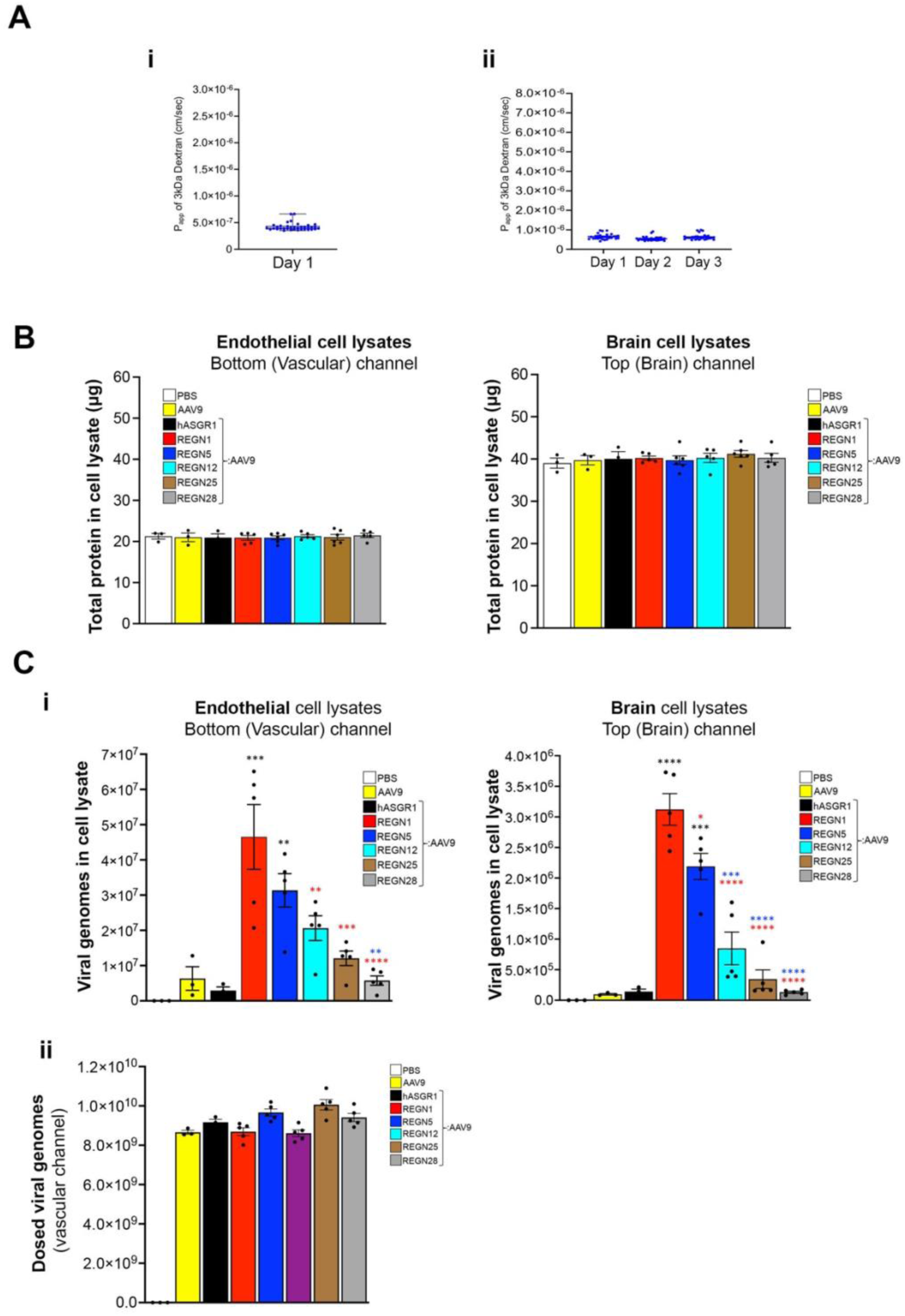
Additional experiments related to the testing of the resolution of the Brain-Chip to detect BBB crossing differences between hTfR1-specific antibodies. (A) Apparent permeability measurements in chips used for testing the hTfR1 antibodies as purified formats (Figure 5) (i) and conjugated to AAV9 (Figure 6) (ii). (**B**) Amount of total protein in endothelial cell and brain cell lysates used for the BBB crossing studies with hTfR1 antibody-conjugated viruses shown in Figure 5. No differences were observed between groups. (C) Quantification of viral genomes in endothelial (i) and brain cells (ii) collected at the end of the AAV9 perfusion (day 3; Figure 6A). The differences between the groups were consistent with the GFP expression analysis experiment (Figure 6) and in agreement with the BBB crossing properties of the antibodies. (ii) Quantification of dosed viral genomes in the vascular media (measurements were performed prior to perfusion). hTfR1:AAV9: N=5 chips for each antibody clone; PBS, AAV9 and hASGR1:AAV9: N=3 chips per group. *p < 0.05, **p < 0.01, ***p < 0.001, ****p< 0.0001, one-way ANOVA with post hoc Tukey’s test. Black asterisks: comparison with PBS, AAV9, hASGR1:AAV9. Red asterisks: comparison with REGN1. Blue asterisks: comparison with REGN5. Averaged data (Mean ± SEM) and individual chip values are shown (A-C).

## REFERENCES

1. Dementia, A. 2021 Alzheimer’s disease facts and figures. Alzheimers Dementia 2021, *17*, 327–406, doi:10.1002/alz.12328.

2. Cai, Y.; Fan, K.; Lin, J.; Ma, L.; Li, F. Advances in BBB on Chip and Application for Studying Reversible Opening of Blood-Brain Barrier by Sonoporation. Micromachines (Basel) 2022, 14, doi:10.3390/mi14010112.

3. Gribkoff, V.K.; Kaczmarek, L.K. The need for new approaches in CNS drug discovery: Why drugs have failed, and what can be done to improve outcomes. Neuropharmacology 2017, 120, 11–19, doi:10.1016/j.neuropharm.2016.03.021.

4. Atkins, J.T.; George, G.C.; Hess, K.; Marcelo-Lewis, K.L.; Yuan, Y.; Borthakur, G.; Khozin, S.; LoRusso, P.; Hong, D.S. Pre-clinical animal models are poor predictors of human toxicities in phase 1 oncology clinical trials. Br J Cancer 2020, 123, 1496–1501, doi:10.1038/s41416-020-01033-x.

5. Van Norman, G.A. Limitations of Animal Studies for Predicting Toxicity in Clinical Trials: Is it Time to Rethink Our Current Approach? JACC Basic Transl Sci 2019, 4, 845–854, doi:10.1016/j.jacbts.2019.10.008.

6. Marshall, L.J.; Bailey, J.; Cassotta, M.; Herrmann, K.; Pistollato, F. Poor Translatability of Biomedical Research Using Animals - A Narrative Review. Altern Lab Anim 2023, 51, 102–135, doi:10.1177/02611929231157756.

7. Loewa, A.; Feng, J.J.; Hedtrich, S. Human disease models in drug development. Nat Rev Bioeng 2023, 1–15, doi:10.1038/s44222-023-00063-3.

8. Sporns, O. The complex brain: connectivity, dynamics, information. Trends Cogn Sci 2022, 26, 1066–1067, doi:10.1016/j.tics.2022.08.002.

9. McConnell, H.L.; Kersch, C.N.; Woltjer, R.L.; Neuwelt, E.A. The Translational Significance of the Neurovascular Unit. J Biol Chem 2017, 292, 762–770, doi:10.1074/jbc.R116.760215.

10. Liebner, S.; Dijkhuizen, R.M.; Reiss, Y.; Plate, K.H.; Agalliu, D.; Constantin, G. Functional morphology of the blood-brain barrier in health and disease. Acta Neuropathol 2018, 135, 311–336, doi:10.1007/s00401-018-1815-1.

11. Kortekaas, R.; Leenders, K.L.; van Oostrom, J.C.; Vaalburg, W.; Bart, J.; Willemsen, A.T.; Hendrikse, N.H. Blood-brain barrier dysfunction in parkinsonian midbrain in vivo. Ann Neurol 2005, 57, 176–179, doi:10.1002/ana.20369.

12. Mayer, M.G.; Fischer, T. Microglia at the blood brain barrier in health and disease. Front Cell Neurosci 2024, 18, 1360195, doi:10.3389/fncel.2024.1360195.

13. Wu, D.; Chen, Q.; Chen, X.; Han, F.; Chen, Z.; Wang, Y. The blood-brain barrier: structure, regulation, and drug delivery. Signal Transduct Target Ther 2023, 8, 217, doi:10.1038/s41392-023-01481-w.

14. Verscheijden, L.F.M.; Koenderink, J.B.; de Wildt, S.N.; Russel, F.G.M. Differences in P-glycoprotein activity in human and rodent blood-brain barrier assessed by mechanistic modelling. Arch Toxicol 2021, 95, 3015–3029, doi:10.1007/s00204-021-03115-y.

15. Wasielewska, J.M.; Da Silva Chaves, J.C.; White, A.R.; Oikari, L.E. Modeling the Blood–Brain Barrier to Understand Drug Delivery in Alzheimer’s Disease. In Alzheimer’s Disease: Drug Discovery, Huang, X., Ed.; Exon Publications

16. Hodge, R.D.; Bakken, T.E.; Miller, J.A.; Smith, K.A.; Barkan, E.R.; Graybuck, L.T.; Close, J.L.; Long, B.; Johansen, N.; Penn, O.;, et al. Conserved cell types with divergent features in human versus mouse cortex. Nature 2019, 573, 61–68, doi:10.1038/s41586-019-1506-7.

17. Urich, E.; Lazic, S.E.; Molnos, J.; Wells, I.; Freskgård, P.O. Transcriptional profiling of human brain endothelial cells reveals key properties crucial for predictive in vitro blood-brain barrier models. PLoS One 2012, 7, e38149, doi:10.1371/journal.pone.0038149.

18. Hoshi, Y.; Uchida, Y.; Tachikawa, M.; Inoue, T.; Ohtsuki, S.; Terasaki, T. Quantitative atlas of blood-brain barrier transporters, receptors, and tight junction proteins in rats and common marmoset. J Pharm Sci 2013, 102, 3343–3355, doi:10.1002/jps.23575.

19. Syvänen, S.; Lindhe, O.; Palner, M.; Kornum, B.R.; Rahman, O.; Långström, B.; Knudsen, G.M.; Hammarlund-Udenaes, M. Species differences in blood-brain barrier transport of three positron emission tomography radioligands with emphasis on P-glycoprotein transport. Drug Metab Dispos 2009, 37, 635–643, doi:10.1124/dmd.108.024745.

20. Oberheim, N.A.; Takano, T.; Han, X.; He, W.; Lin, J.H.; Wang, F.; Xu, Q.; Wyatt, J.D.; Pilcher, W.; Ojemann, J.G.;, et al. Uniquely hominid features of adult human astrocytes. J Neurosci 2009, 29, 3276–3287, doi:10.1523/jneurosci.4707-08.2009.

21. Eidsvaag, V.A.; Enger, R.; Hansson, H.A.; Eide, P.K.; Nagelhus, E.A. Human and mouse cortical astrocytes differ in aquaporin-4 polarization toward microvessels. Glia 2017, 65, 964–973, doi:10.1002/glia.23138.

22. Zeppenfeld, D.M.; Simon, M.; Haswell, J.D.; D’Abreo, D.; Murchison, C.; Quinn, J.F.; Grafe, M.R.; Woltjer, R.L.; Kaye, J.; Iliff, J.J. Association of Perivascular Localization of Aquaporin-4 With Cognition and Alzheimer Disease in Aging Brains. JAMA Neurol 2017, 74, 91–99, doi:10.1001/jamaneurol.2016.4370.

23. Terstappen, G.C.; Meyer, A.H.; Bell, R.D.; Zhang, W. Strategies for delivering therapeutics across the blood-brain barrier. Nat Rev Drug Discov 2021, 20, 362–383, doi:10.1038/s41573-021-00139-y.

24. Bhunia, S.; Kolishetti, N.; Vashist, A.; Yndart Arias, A.; Brooks, D.; Nair, M. Drug Delivery to the Brain: Recent Advances and Unmet Challenges. Pharmaceutics 2023, 15, doi:10.3390/pharmaceutics15122658.

25. Achar, A.; Myers, R.; Ghosh, C. Drug Delivery Challenges in Brain Disorders across the Blood-Brain Barrier: Novel Methods and Future Considerations for Improved Therapy. Biomedicines 2021, 9, doi:10.3390/biomedicines9121834.

26. Stephenson, D.; Belfiore-Oshan, R.; Karten, Y.; Keavney, J.; Kwok, D.K.; Martinez, T.; Montminy, J.; Müller, M.; Romero, K.; Sivakumaran, S. Transforming Drug Development for Neurological Disorders: Proceedings from a Multidisease Area Workshop. Neurotherapeutics 2023, 20, 1682–1691, doi:10.1007/s13311-023-01440-x.

27. Schneider, L. FDA no longerhas to requireanimal testingfor new drugs. Science 2023, 379 127–128.

28. Ingber, D.E. Human organs-on-chips for disease modelling, drug development and personalized medicine. Nat Rev Genet 2022, 23, 467–491, doi:10.1038/s41576-022-00466-9.

29. Bhatia, S.N.; Ingber, D.E. Microfluidic organs-on-chips. Nat Biotechnol 2014, 32, 760–772, doi:10.1038/nbt.2989.

30. Haring, A.P.; Sontheimer, H.; Johnson, B.N. Microphysiological Human Brain and Neural Systems-on-a-Chip: Potential Alternatives to Small Animal Models and Emerging Platforms for Drug Discovery and Personalized Medicine. Stem Cell Rev Rep 2017, 13, 381–406, doi:10.1007/s12015-017-9738-0.

31. Mulay, A.R.; Hwang, J.; Kim, D.H. Microphysiological Blood-Brain Barrier Systems for Disease Modeling and Drug Development. Adv Healthc Mater 2024, e2303180, doi:10.1002/adhm.202303180.

32. Guarino, V.; Zizzari, A.; Bianco, M.; Gigli, G.; Moroni, L.; Arima, V. Advancements in modelling human blood brain-barrier on a chip. Biofabrication 2023, 15, doi:10.1088/1758-5090/acb571.

33. Jagadeesan, S.; Workman, M.J.; Herland, A.; Svendsen, C.N.; Vatine, G.D. Generation of a Human iPSC-Based Blood-Brain Barrier Chip. J Vis Exp 2020, doi:10.3791/60925.

34. Vatine, G.D.; Barrile, R.; Workman, M.J.; Sances, S.; Barriga, B.K.; Rahnama, M.; Barthakur, S.; Kasendra, M.; Lucchesi, C.; Kerns, J.;, et al. Human iPSC-Derived Blood-Brain Barrier Chips Enable Disease Modeling and Personalized Medicine Applications. Cell Stem Cell 2019, 24, 995–1005.e1006, doi:10.1016/j.stem.2019.05.011.

35. Choi, J.H.; Santhosh, M.; Choi, J.W. In Vitro Blood-Brain Barrier-Integrated Neurological Disorder Models Using a Microfluidic Device. Micromachines (Basel) 2019, 11, doi:10.3390/mi11010021.

36. Yang, J.Y.; Shin, D.S.; Jeong, M.; Kim, S.S.; Jeong, H.N.; Lee, B.H.; Hwang, K.S.; Son, Y.; Jeong, H.C.; Choi, C.H.;, et al. Evaluation of Drug Blood-Brain-Barrier Permeability Using a Microfluidic Chip. Pharmaceutics 2024, 16, doi:10.3390/pharmaceutics16050574.

37. Choi, J.W.; Kim, K.; Mukhambetiyar, K.; Lee, N.K.; Sabaté Del Río, J.; Joo, J.; Park, C.G.; Kwon, T.; Park, T.E. Organ-on-a-Chip Approach for Accelerating Blood-Brain Barrier Nanoshuttle Discovery. ACS Nano 2024, 18, 14388–14402, doi:10.1021/acsnano.4c00994.

38. Burgio, F.; Gaiser, C.; Brady, K.; Gatta, V.; Class, R.; Schrage, R.; Suter-Dick, L. A Perfused In Vitro Human iPSC-Derived Blood-Brain Barrier Faithfully Mimics Transferrin Receptor-Mediated Transcytosis of Therapeutic Antibodies. Cell Mol Neurobiol 2023, 43, 4173–4187, doi:10.1007/s10571-023-01404-x.

39. Kaplan, L.; Chow, B.W.; Gu, C. Neuronal regulation of the blood-brain barrier and neurovascular coupling. Nat Rev Neurosci 2020, 21, 416–432, doi:10.1038/s41583-020-0322-2.

40. Pediaditakis, I.; Kodella, K.R.; Manatakis, D.V.; Le, C.Y.; Barthakur, S.; Sorets, A.; Gravanis, A.; Ewart, L.; Rubin, L.L.; Manolakos, E.S.;, et al. A microengineered Brain-Chip to model neuroinflammation in humans. iScience 2022, 25, 104813, doi:10.1016/j.isci.2022.104813.

41. Pediaditakis, I.; Kodella, K.R.; Manatakis, D.V.; Le, C.Y.; Hinojosa, C.D.; Tien-Street, W.; Manolakos, E.S.; Vekrellis, K.; Hamilton, G.A.; Ewart, L.;, et al. Modeling alpha-synuclein pathology in a human brain-chip to assess blood-brain barrier disruption. Nat Commun 2021, 12, 5907, doi:10.1038/s41467-021-26066-5.

42. de Rus Jacquet, A.; Alpaugh, M.; Denis, H.L.; Tancredi, J.L.; Boutin, M.; Decaestecker, J.; Beauparlant, C.; Herrmann, L.; Saint-Pierre, M.; Parent, M.;, et al. The contribution of inflammatory astrocytes to BBB impairments in a brain-chip model of Parkinson’s disease. Nat Commun 2023, 14, 3651, doi:10.1038/s41467-023-39038-8.

43. Nakaso, K. Roles of Microglia in Neurodegenerative Diseases. Yonago Acta Med 2024, 67, 1–8, doi:10.33160/yam.2024.02.001.

44. Gao, C.; Jiang, J.; Tan, Y.; Chen, S. Microglia in neurodegenerative diseases: mechanism and potential therapeutic targets. Signal Transduct Target Ther 2023, 8, 359, doi:10.1038/s41392-023-01588-0.

45. Hickman, S.; Izzy, S.; Sen, P.; Morsett, L.; El Khoury, J. Microglia in neurodegeneration. Nat Neurosci 2018, 21, 1359–1369, doi:10.1038/s41593-018-0242-x.

46. Muzio, L.; Viotti, A.; Martino, G. Microglia in Neuroinflammation and Neurodegeneration: From Understanding to Therapy. Front Neurosci 2021, 15, 742065, doi:10.3389/fnins.2021.742065.

47. Lee, J.W.; Chun, W.; Lee, H.J.; Kim, S.M.; Min, J.H.; Kim, D.Y.; Kim, M.O.; Ryu, H.W.; Lee, S.U. The Role of Microglia in the Development of Neurodegenerative Diseases. Biomedicines 2021, 9, doi:10.3390/biomedicines9101449.

48. Alvarez, J.I.; Katayama, T.; Prat, A. Glial influence on the blood brain barrier. Glia 2013, 61, 1939–1958, doi:10.1002/glia.22575.

49. Yuan, W.; Lv, Y.; Zeng, M.; Fu, B.M. Non-invasive measurement of solute permeability in cerebral microvessels of the rat. Microvasc Res 2009, 77, 166–173, doi:10.1016/j.mvr.2008.08.004.

50. Shi, L.; Zeng, M.; Sun, Y.; Fu, B.M. Quantification of blood-brain barrier solute permeability and brain transport by multiphoton microscopy. J Biomech Eng 2014, 136, 031005, doi:10.1115/1.4025892.

51. Khalilov, I.; Khazipov, R.; Esclapez, M.; Ben-Ari, Y. Bicuculline induces ictal seizures in the intact hippocampus recorded in vitro. Eur J Pharmacol 1997, 319, R5–6, doi:10.1016/s0014-2999(96)00964-8.

52. Xu, A.; Cui, S.; Wang, J.H. Incoordination among Subcellular Compartments Is Associated with Depression-Like Behavior Induced by Chronic Mild Stress. Int J Neuropsychopharmacol 2016, 19, doi:10.1093/ijnp/pyv122.

53. Samoilova, M.; Li, J.; Pelletier, M.R.; Wentlandt, K.; Adamchik, Y.; Naus, C.C.; Carlen, P.L. Epileptiform activity in hippocampal slice cultures exposed chronically to bicuculline: increased gap junctional function and expression. J Neurochem 2003, 86, 687–699, doi:10.1046/j.1471-4159.2003.01893.x.

54. Peng, B.; Tong, Z.; Tong, W.Y.; Pasic, P.J.; Oddo, A.; Dai, Y.; Luo, M.; Frescene, J.; Welch, N.G.; Easton, C.D.;, et al. In Situ Surface Modification of Microfluidic Blood-Brain-Barriers for Improved Screening of Small Molecules and Nanoparticles. ACS Appl Mater Interfaces 2020, 12, 56753–56766, doi:10.1021/acsami.0c17102.

55. Rochfort, K.D.; Collins, L.E.; Murphy, R.P.; Cummins, P.M. Downregulation of blood-brain barrier phenotype by proinflammatory cytokines involves NADPH oxidase-dependent ROS generation: consequences for interendothelial adherens and tight junctions. PLoS One 2014, 9, e101815, doi:10.1371/journal.pone.0101815.

56. Wei, W.J.; Wang, Y.C.; Guan, X.; Chen, W.G.; Liu, J. A neurovascular unit-on-a-chip: culture and differentiation of human neural stem cells in a three-dimensional microfluidic environment. Neural Regen Res 2022, 17, 2260–2266, doi:10.4103/1673-5374.337050.

57. Pan, H.; Li, H.; Guo, S.; Wang, C.; Long, L.; Wang, X.; Shi, H.; Zhang, K.; Chen, H.; Li, S. The mechanisms and functions of TNF-α in intervertebral disc degeneration. Exp Gerontol 2023, 174, 112119, doi:10.1016/j.exger.2023.112119.

58. Kouchaki, E.; Kakhaki, R.D.; Tamtaji, O.R.; Dadgostar, E.; Behnam, M.; Nikoueinejad, H.; Akbari, H. Increased serum levels of TNF-α and decreased serum levels of IL-27 in patients with Parkinson disease and their correlation with disease severity. Clin Neurol Neurosurg 2018, 166, 76–79, doi:10.1016/j.clineuro.2018.01.022.

59. Probert, L. TNF and its receptors in the CNS: The essential, the desirable and the deleterious effects. Neuroscience 2015, 302, 2–22, doi:10.1016/j.neuroscience.2015.06.038.

60. Versele, R.; Sevin, E.; Gosselet, F.; Fenart, L.; Candela, P. TNF-α and IL-1β Modulate Blood-Brain Barrier Permeability and Decrease Amyloid-β Peptide Efflux in a Human Blood-Brain Barrier Model. Int J Mol Sci 2022, 23, doi:10.3390/ijms231810235.

61. Chen, A.Q.; Fang, Z.; Chen, X.L.; Yang, S.; Zhou, Y.F.; Mao, L.; Xia, Y.P.; Jin, H.J.; Li, Y.N.; You, M.F.;, et al. Microglia-derived TNF-α mediates endothelial necroptosis aggravating blood brain-barrier disruption after ischemic stroke. Cell Death Dis 2019, 10, 487, doi:10.1038/s41419-019-1716-9.

62. Cheng, Y.; Desse, S.; Martinez, A.; Worthen, R.J.; Jope, R.S.; Beurel, E. TNFα disrupts blood brain barrier integrity to maintain prolonged depressive-like behavior in mice. Brain Behav Immun 2018, 69, 556–567, doi:10.1016/j.bbi.2018.02.003.

63. Schenk, G.J.; de Vries, H.E. Altered blood-brain barrier transport in neuro-inflammatory disorders. Drug Discov Today Technol 2016, 20, 5–11, doi:10.1016/j.ddtec.2016.07.002.

64. Obermeier, B.; Daneman, R.; Ransohoff, R.M. Development, maintenance and disruption of the blood-brain barrier. Nat Med 2013, 19, 1584–1596, doi:10.1038/nm.3407.

65. Shim, H.G.; Jang, S.S.; Kim, S.H.; Hwang, E.M.; Min, J.O.; Kim, H.Y.; Kim, Y.S.; Ryu, C.; Chung, G.; Kim, Y.;, et al. TNF-α increases the intrinsic excitability of cerebellar Purkinje cells through elevating glutamate release in Bergmann Glia. Sci Rep 2018, 8, 11589, doi:10.1038/s41598-018-29786-9.

66. Clark, I.A.; Vissel, B. Excess cerebral TNF causing glutamate excitotoxicity rationalizes treatment of neurodegenerative diseases and neurogenic pain by anti-TNF agents. J Neuroinflammation 2016, 13, 236, doi:10.1186/s12974-016-0708-2.

67. Olmos, G.; Lladó, J. Tumor necrosis factor alpha: a link between neuroinflammation and excitotoxicity. Mediators Inflamm 2014, 2014, 861231, doi:10.1155/2014/861231.

68. Ye, L.; Huang, Y.; Zhao, L.; Li, Y.; Sun, L.; Zhou, Y.; Qian, G.; Zheng, J.C. IL-1β and TNF-α induce neurotoxicity through glutamate production: a potential role for neuronal glutaminase. J Neurochem 2013, 125, 897–908, doi:10.1111/jnc.12263.

69. Rochfort, K.D.; Cummins, P.M. The blood-brain barrier endothelium: a target for pro-inflammatory cytokines. Biochem Soc Trans 2015, 43, 702–706, doi:10.1042/bst20140319.

70. Nishioku, T.; Matsumoto, J.; Dohgu, S.; Sumi, N.; Miyao, K.; Takata, F.; Shuto, H.; Yamauchi, A.; Kataoka, Y. Tumor necrosis factor-alpha mediates the blood-brain barrier dysfunction induced by activated microglia in mouse brain microvascular endothelial cells. J Pharmacol Sci 2010, 112, 251–254, doi:10.1254/jphs.09292sc.

71. Poller, B.; Drewe, J.; Krähenbühl, S.; Huwyler, J.; Gutmann, H. Regulation of BCRP (ABCG2) and P-glycoprotein (ABCB1) by cytokines in a model of the human blood-brain barrier. Cell Mol Neurobiol 2010, 30, 63–70, doi:10.1007/s10571-009-9431-1.

72. Shao, F.; Wang, X.; Wu, H.; Wu, Q.; Zhang, J. Microglia and Neuroinflammation: Crucial Pathological Mechanisms in Traumatic Brain Injury-Induced Neurodegeneration. Front Aging Neurosci 2022, 14, 825086, doi:10.3389/fnagi.2022.825086.

73. Gullotta, G.S.; Costantino, G.; Sortino, M.A.; Spampinato, S.F. Microglia and the Blood-Brain Barrier: An External Player in Acute and Chronic Neuroinflammatory Conditions. Int J Mol Sci 2023, 24, doi:10.3390/ijms24119144.

74. Takata, F.; Nakagawa, S.; Matsumoto, J.; Dohgu, S. Blood-Brain Barrier Dysfunction Amplifies the Development of Neuroinflammation: Understanding of Cellular Events in Brain Microvascular Endothelial Cells for Prevention and Treatment of BBB Dysfunction. Front Cell Neurosci 2021, 15, 661838, doi:10.3389/fncel.2021.661838.

75. Haruwaka, K.; Ikegami, A.; Tachibana, Y.; Ohno, N.; Konishi, H.; Hashimoto, A.; Matsumoto, M.; Kato, D.; Ono, R.; Kiyama, H.;, et al. Dual microglia effects on blood brain barrier permeability induced by systemic inflammation. Nat Commun 2019, 10, 5816, doi:10.1038/s41467-019-13812-z.

76. Laffer, B.; Bauer, D.; Wasmuth, S.; Busch, M.; Jalilvand, T.V.; Thanos, S.; Meyer Zu Hörste, G.; Loser, K.; Langmann, T.; Heiligenhaus, A.; et al. Loss of IL-10 Promotes Differentiation of Microglia to a M1 Phenotype. Front Cell Neurosci 2019, 13, 430, doi:10.3389/fncel.2019.00430.

77. Johnsen, K.B.; Burkhart, A.; Melander, F.; Kempen, P.J.; Vejlebo, J.B.; Siupka, P.; Nielsen, M.S.; Andresen, T.L.; Moos, T. Targeting transferrin receptors at the blood-brain barrier improves the uptake of immunoliposomes and subsequent cargo transport into the brain parenchyma. Sci Rep 2017, 7, 10396, doi:10.1038/s41598-017-11220-1.

78. Sonoda, H.; Morimoto, H.; Yoden, E.; Koshimura, Y.; Kinoshita, M.; Golovina, G.; Takagi, H.; Yamamoto, R.; Minami, K.; Mizoguchi, A.;, et al. A Blood-Brain-Barrier-Penetrating Anti-human Transferrin Receptor Antibody Fusion Protein for Neuronopathic Mucopolysaccharidosis II. Mol Ther 2018, 26, 1366–1374, doi:10.1016/j.ymthe.2018.02.032.

79. Lee, H.J.; Engelhardt, B.; Lesley, J.; Bickel, U.; Pardridge, W.M. Targeting rat anti-mouse transferrin receptor monoclonal antibodies through blood-brain barrier in mouse. J Pharmacol Exp Ther 2000, 292, 1048–1052.

80. Dawson, T.M.; Golde, T.E.; Lagier-Tourenne, C. Animal models of neurodegenerative diseases. Nat Neurosci 2018, 21, 1370–1379, doi:10.1038/s41593-018-0236-8.

81. Kumar, V.; Sanseau, P.; Simola, D.F.; Hurle, M.R.; Agarwal, P. Systematic Analysis of Drug Targets Confirms Expression in Disease-Relevant Tissues. Sci Rep 2016, 6, 36205, doi:10.1038/srep36205.

82. Cardoso-Moreira, M.; Sarropoulos, I.; Velten, B.; Mort, M.; Cooper, D.N.; Huber, W.; Kaessmann, H. Developmental Gene Expression Differences between Humans and Mammalian Models. Cell Rep 2020, 33, 108308, doi:10.1016/j.celrep.2020.108308.

83. Franzen, N.; van Harten, W.H.; Retèl, V.P.; Loskill, P.; van den Eijnden-van Raaij, J.; M, I.J. Impact of organ-on-a-chip technology on pharmaceutical R&D costs. Drug Discov Today 2019, 24, 1720–1724, doi:10.1016/j.drudis.2019.06.003.

84. Farhang Doost, N.; Srivastava, S.K. A Comprehensive Review of Organ-on-a-Chip Technology and Its Applications. Biosensors (Basel) 2024, 14, doi:10.3390/bios14050225.

85. Clayton, K.A.; Van Enoo, A.A.; Ikezu, T. Alzheimer’s Disease: The Role of Microglia in Brain Homeostasis and Proteopathy. Front Neurosci 2017, 11, 680, doi:10.3389/fnins.2017.00680.

86. Rodríguez-Arellano, J.J.; Parpura, V.; Zorec, R.; Verkhratsky, A. Astrocytes in physiological aging and Alzheimer’s disease. Neuroscience 2016, 323, 170–182, doi:10.1016/j.neuroscience.2015.01.007.

87. Cai, Z.; Xiao, M. Oligodendrocytes and Alzheimer’s disease. Int J Neurosci 2016, 126, 97–104, doi:10.3109/00207454.2015.1025778.

88. Marsan, E.; Velmeshev, D.; Ramsey, A.; Patel, R.K.; Zhang, J.; Koontz, M.; Andrews, M.G.; de Majo, M.; Mora, C.; Blumenfeld, J.;, et al. Astroglial toxicity promotes synaptic degeneration in the thalamocortical circuit in frontotemporal dementia with GRN mutations. J Clin Invest 2023, 133, doi:10.1172/jci164919.

89. Brandebura, A.N.; Paumier, A.; Onur, T.S.; Allen, N.J. Astrocyte contribution to dysfunction, risk and progression in neurodegenerative disorders. Nat Rev Neurosci 2023, 24, 23–39, doi:10.1038/s41583-022-00641-1.

90. Han, S.; Gim, Y.; Jang, E.H.; Hur, E.M. Functions and dysfunctions of oligodendrocytes in neurodegenerative diseases. Front Cell Neurosci 2022, 16, 1083159, doi:10.3389/fncel.2022.1083159.

91. Yuan, Y.; Sun, J.; Dong, Q.; Cui, M. Blood-brain barrier endothelial cells in neurodegenerative diseases: Signals from the “barrier”. Front Neurosci 2023, 17, 1047778, doi:10.3389/fnins.2023.1047778.

92. Fang, Y.C.; Hsieh, Y.C.; Hu, C.J.; Tu, Y.K. Endothelial Dysfunction in Neurodegenerative Diseases. Int J Mol Sci 2023, 24, doi:10.3390/ijms24032909.

93. Miedema, S.S.M.; Mol, M.O.; Koopmans, F.T.W.; Hondius, D.C.; van Nierop, P.; Menden, K.; de Veij Mestdagh, C.F.; van Rooij, J.; Ganz, A.B.; Paliukhovich, I.;, et al. Distinct cell type-specific protein signatures in GRN and MAPT genetic subtypes of frontotemporal dementia. Acta Neuropathol Commun 2022, 10, 100, doi:10.1186/s40478-022-01387-8.

94. Bright, F.; Werry, E.L.; Dobson-Stone, C.; Piguet, O.; Ittner, L.M.; Halliday, G.M.; Hodges, J.R.; Kiernan, M.C.; Loy, C.T.; Kassiou, M.;, et al. Neuroinflammation in frontotemporal dementia. Nat Rev Neurol 2019, 15, 540–555, doi:10.1038/s41582-019-0231-z.

95. Barreras, P.; Pamies, D.; Hartung, T.; Pardo, C.A. Human brain microphysiological systems in the study of neuroinfectious disorders. Exp Neurol 2023, 365, 114409, doi:10.1016/j.expneurol.2023.114409.

96. Boylin, K.; Aquino, G.V.; Purdon, M.; Abedi, K.; Kasendra, M.; Barrile, R. Basic models to advanced systems: harnessing the power of organoids-based microphysiological models of the human brain. Biofabrication 2024, 16, doi:10.1088/1758-5090/ad4c08.

97. Bai, W.; Zhou, Y.G. Homeostasis of the Intraparenchymal-Blood Glutamate Concentration Gradient: Maintenance, Imbalance, and Regulation. Front Mol Neurosci 2017, 10, 400, doi:10.3389/fnmol.2017.00400.

98. Hutchinson, P.J.; O’Connell, M.T.; Al-Rawi, P.G.; Kett-White, C.R.; Gupta, A.K.; Maskell, L.B.; Pickard, J.D.; Kirkpatrick, P.J. Increases in GABA concentrations during cerebral ischaemia: a microdialysis study of extracellular amino acids. J Neurol Neurosurg Psychiatry 2002, 72, 99–105, doi:10.1136/jnnp.72.1.99.

99. Shen, H.; Kihara, T.; Hongo, H.; Wu, X.; Kem, W.R.; Shimohama, S.; Akaike, A.; Niidome, T.; Sugimoto, H. Neuroprotection by donepezil against glutamate excitotoxicity involves stimulation of alpha7 nicotinic receptors and internalization of NMDA receptors. Br J Pharmacol 2010, 161, 127–139, doi:10.1111/j.1476-5381.2010.00894.x.

100. Simões, A.P.; Silva, C.G.; Marques, J.M.; Pochmann, D.; Porciúncula, L.O.; Ferreira, S.; Oses, J.P.; Beleza, R.O.; Real, J.I.; Köfalvi, A.;, et al. Glutamate-induced and NMDA receptor-mediated neurodegeneration entails P2Y1 receptor activation. Cell Death Dis 2018, 9, 297, doi:10.1038/s41419-018-0351-1.

101. Xiong, Z.Q.; McNamara, J.O. Fleeting activation of ionotropic glutamate receptors sensitizes cortical neurons to complement attack. Neuron 2002, 36, 363–374, doi:10.1016/s0896-6273(02)00977-7.

102. Ghatak, S.; Talantova, M.; McKercher, S.R.; Lipton, S.A. Novel Therapeutic Approach for Excitatory/Inhibitory Imbalance in Neurodevelopmental and Neurodegenerative Diseases. Annu Rev Pharmacol Toxicol 2021, 61, 701–721, doi:10.1146/annurev-pharmtox-032320-015420.

103. Gao, R.; Penzes, P. Common mechanisms of excitatory and inhibitory imbalance in schizophrenia and autism spectrum disorders. Curr Mol Med 2015, 15, 146–167, doi:10.2174/1566524015666150303003028.

104. van van Hugte, E.J.H.; Schubert, D.; Nadif Kasri, N. Excitatory/inhibitory balance in epilepsies and neurodevelopmental disorders: Depolarizing γ-aminobutyric acid as a common mechanism. Epilepsia 2023, 64, 1975–1990, doi:10.1111/epi.17651.

105. Frankola, K.A.; Greig, N.H.; Luo, W.; Tweedie, D. Targeting TNF-α to elucidate and ameliorate neuroinflammation in neurodegenerative diseases. CNS Neurol Disord Drug Targets 2011, 10, 391–403, doi:10.2174/187152711794653751.

106. Amin, R.; Quispe, C.; Docea, A.O.; Ydyrys, A.; Kulbayeva, M.; Durna Daştan, S.; Calina, D.; Sharifi-Rad, J. The role of Tumour Necrosis Factor in neuroinflammation associated with Parkinson’s disease and targeted therapies. Neurochem Int 2022, 158, 105376, doi:10.1016/j.neuint.2022.105376.

107. Ou, W.; Yang, J.; Simanauskaite, J.; Choi, M.; Castellanos, D.M.; Chang, R.; Sun, J.; Jagadeesan, N.; Parfitt, K.D.; Cribbs, D.H.;, et al. Biologic TNF-α inhibitors reduce microgliosis, neuronal loss, and tau phosphorylation in a transgenic mouse model of tauopathy. J Neuroinflammation 2021, 18, 312, doi:10.1186/s12974-021-02332-7.

108. Henning, L.; Antony, H.; Breuer, A.; Müller, J.; Seifert, G.; Audinat, E.; Singh, P.; Brosseron, F.; Heneka, M.T.; Steinhäuser, C.;, et al. Reactive microglia are the major source of tumor necrosis factor alpha and contribute to astrocyte dysfunction and acute seizures in experimental temporal lobe epilepsy. Glia 2023, 71, 168–186, doi:10.1002/glia.24265.

109. Patel, D.C.; Wallis, G.; Dahle, E.J.; McElroy, P.B.; Thomson, K.E.; Tesi, R.J.; Szymkowski, D.E.; West, P.J.; Smeal, R.M.; Patel, M.;, et al. Hippocampal TNFα Signaling Contributes to Seizure Generation in an Infection-Induced Mouse Model of Limbic Epilepsy. eNeuro 2017, 4, doi:10.1523/eneuro.0105-17.2017.

110. Zhang, W.; Xiao, D.; Mao, Q.; Xia, H. Role of neuroinflammation in neurodegeneration development. Signal Transduct Target Ther 2023, 8, 267, doi:10.1038/s41392-023-01486-5.

111. Li, W.; Wu, J.; Zeng, Y.; Zheng, W. Neuroinflammation in epileptogenesis: from pathophysiology to therapeutic strategies. Front Immunol 2023, 14, 1269241, doi:10.3389/fimmu.2023.1269241.

112. Cereda, C.; Baiocchi, C.; Bongioanni, P.; Cova, E.; Guareschi, S.; Metelli, M.R.; Rossi, B.; Sbalsi, I.; Cuccia, M.C.; Ceroni, M. TNF and sTNFR1/2 plasma levels in ALS patients. J Neuroimmunol 2008, 194, 123–131, doi:10.1016/j.jneuroim.2007.10.028.

113. Jones, A.R.; Shusta, E.V. Blood-brain barrier transport of therapeutics via receptor-mediation. Pharm Res 2007, 24, 1759–1771, doi:10.1007/s11095-007-9379-0.

114. Pardridge, W.M. Advanced Blood-Brain Barrier Drug Delivery. Pharmaceutics 2022, 15, doi:10.3390/pharmaceutics15010093.

115. Zhang, X.; He, T.; Chai, Z.; Samulski, R.J.; Li, C. Blood-brain barrier shuttle peptides enhance AAV transduction in the brain after systemic administration. Biomaterials 2018, 176, 71–83, doi:10.1016/j.biomaterials.2018.05.041.

116. Huang, Q.; Chan, K.Y.; Wu, J.; Botticello-Romero, N.R.; Zheng, Q.; Lou, S.; Keyes, C.; Svanbergsson, A.; Johnston, J.; Mills, A.;, et al. An AAV capsid reprogrammed to bind human transferrin receptor mediates brain-wide gene delivery. Science 2024, 384, 1220–1227, doi:10.1126/science.adm8386.

117. Lajoie, J.M.; Shusta, E.V. Targeting receptor-mediated transport for delivery of biologics across the blood-brain barrier. Annu Rev Pharmacol Toxicol 2015, 55, 613–631, doi:10.1146/annurev-pharmtox-010814-124852.

118. Pulgar, V.M. Transcytosis to Cross the Blood Brain Barrier, New Advancements and Challenges. Front Neurosci 2018, 12, 1019, doi:10.3389/fnins.2018.01019.

119. Thomsen, M.S.; Johnsen, K.B.; Kucharz, K.; Lauritzen, M.; Moos, T. Blood-Brain Barrier Transport of Transferrin Receptor-Targeted Nanoparticles. Pharmaceutics 2022, 14, doi:10.3390/pharmaceutics14102237.

120. Johnsen, K.B.; Burkhart, A.; Thomsen, L.B.; Andresen, T.L.; Moos, T. Targeting the transferrin receptor for brain drug delivery. Prog Neurobiol 2019, 181, 101665, doi:10.1016/j.pneurobio.2019.101665.

121. Niewoehner, J.; Bohrmann, B.; Collin, L.; Urich, E.; Sade, H.; Maier, P.; Rueger, P.; Stracke, J.O.; Lau, W.; Tissot, A.C.;, et al. Increased brain penetration and potency of a therapeutic antibody using a monovalent molecular shuttle. Neuron 2014, 81, 49–60, doi:10.1016/j.neuron.2013.10.061.

122. Curzer, H.J.; Perry, G.; Wallace, M.C.; Perry, D. The Three Rs of Animal Research: What they Mean for the Institutional Animal Care and Use Committee and Why. Sci Eng Ethics 2016, 22, 549–565, doi:10.1007/s11948-015-9659-8.

123. Lu, T.M.; Houghton, S.; Magdeldin, T.; Durán, J.G.B.; Minotti, A.P.; Snead, A.; Sproul, A.; Nguyen, D.T.; Xiang, J.; Fine, H.A.;, et al. Pluripotent stem cell-derived epithelium misidentified as brain microvascular endothelium requires ETS factors to acquire vascular fate. Proc Natl Acad Sci U S A 2021, 118, doi:10.1073/pnas.2016950118.

124. Qian, T.; Maguire, S.E.; Canfield, S.G.; Bao, X.; Olson, W.R.; Shusta, E.V.; Palecek, S.P. Directed differentiation of human pluripotent stem cells to blood-brain barrier endothelial cells. Sci Adv 2017, 3, e1701679, doi:10.1126/sciadv.1701679.

125. Deli, M.A.; Abrahám, C.S.; Kataoka, Y.; Niwa, M. Permeability studies on in vitro blood-brain barrier models: physiology, pathology, and pharmacology. Cell Mol Neurobiol 2005, 25, 59–127, doi:10.1007/s10571-004-1377-8.

126. Benson, K.; Cramer, S.; Galla, H.J. Impedance-based cell monitoring: barrier properties and beyond. Fluids Barriers CNS 2013, 10, 5, doi:10.1186/2045-8118-10-5.

127. Park, T.E.; Mustafaoglu, N.; Herland, A.; Hasselkus, R.; Mannix, R.; FitzGerald, E.A.; Prantil-Baun, R.; Watters, A.; Henry, O.; Benz, M.;, et al. Hypoxia-enhanced Blood-Brain Barrier Chip recapitulates human barrier function and shuttling of drugs and antibodies. Nat Commun 2019, 10, 2621, doi:10.1038/s41467-019-10588-0.

128. Baker, T.K.; Van Vleet, T.R.; Mahalingaiah, P.K.; Grandhi, T.S.P.; Evers, R.; Ekert, J.; Gosset, J.R.; Chacko, S.A.; Kopec, A.K. The Current Status and Use of Microphysiological Systems by the Pharmaceutical Industry: The International Consortium for Innovation and Quality Microphysiological Systems Affiliate Survey and Commentary. Drug Metab Dispos 2024, 52, 198–209, doi:10.1124/dmd.123.001510.

